# Pathogenic measles viruses cannot evolve to bypass vaccine-induced neutralizing antibodies

**DOI:** 10.1101/2020.10.22.351189

**Authors:** Miguel Ángel Muñoz-Alía, Rebecca A. Nace, Lianwen Zhang, Stephen J. Russell

## Abstract

After centuries of pestilence and decades of global vaccination, measles serotypes capable of evading vaccine-induced immunity have not emerged. Here, by systematically building mutations into the H-glycoprotein of an attenuated measles strain and assaying for serum neutralization, we show that virus evolution is severely constrained by the existence of numerous codominant H-glycoprotein antigenic sites, some critical for binding to pathogenicity receptors SLAMF1 and Nectin-4. We further demonstrate the existence in serum of protective neutralizing antibodies targeting codominant F-glycoprotein epitopes. Calculations suggest that evolution of pathogenic measles viruses capable of escaping serum neutralization in vaccinated individuals is a near-zero probability scenario.

---

Measles (MeV) is a highly transmissible airborne pathogen that spreads systemically and causes transient immune suppression (*1, 2*). Childhood infection is associated with significant mortality and MeV elimination remains a high priority for the World Health Organization (*3*). MeV cell entry via the immune cell receptor SLAMF1 (CD150) drives MeV immunopathogenesis (*4*), whereas a second epithelial cell receptor, NECTIN-4 (PVRL4) is exploited for virus transmission (*5*). Receptor attachment and virus entry are mediated by the concerted action of the hemagglutinin (H) and fusion (F) surface glycoproteins (*2*). The MeV polymerase has a high mutation rate and a correspondingly high frequency of monoclonal nAb-escape mutants but has nevertheless remained monotypic (*6, 7*). Vaccination with a lab-adapted isolate of the genotype A strain MeV, isolated from the throat of David Edmonston in 1954, still confers full protection against all currently circulating genotypes. The evolutionary stability of MeV remains poorly understood but could be due to the inability of its surface glycoproteins to tolerate sequence modification (*8–10*), the multiplicity of B cell epitopes displayed on its surface and/or the low likelihood of mutational escape from combined B and T cell mediated antiviral defences.

Here, to elucidate the serotypic constraints on MeV evolution, we introduced mutated hemagglutinin glycoproteins into the Moraten MeV vaccine whose attenuated phenotype is highly stable with no recorded cases of reversion to pathogenicity or person to person transmission (*11*). Mechanistically, vaccine attenuation is multifactorial, arising from the acquisition of CD46 tropism (*12*), inactivating mutations in V and C immune combat proteins, mutations in the L (polymerase) protein (*13*), and mutations in noncoding sequences (*14*).

To facilitate the detailed analysis of MeV-H neutralization we assembled a comprehensive set of 30 published anti-MeV-H monoclonal antibodies known to neutralize virus infectivity. For those antibodies with unknown target epitopes we propagated the virus in the presence of the antibody, selected escape mutants and sequenced them for epitope identification. Subsequently, we introduced increasing numbers and varying combinations of epitope escape mutations into an antigenically advanced and relatively malleable MeV-H protein obtained from the H1 genotype of MeV (*15*) which, to facilitate the evaluation of mutants with disrupted SLAMF1 and NECTIN-4 receptor binding sites, had been engineered to bind to CD46 by introducing mutations N481Y, H495R, and S546G (**Fig. S1**). We thereby generated a CD46-tropic MeV (MeV-H Δ8) in which all 7 previously reported antigenic sites (*15*), plus an 8^th^ site (IIc) identified in the course of the current study were successfully disrupted **(Fig. S2 and Fig. S3).** Neutralization assays confirmed that viruses encoding the MeV-H Δ8 showed a decreased susceptibility to neutralization by all 30 mAbs **(Fig. 1A and Fig. S4)**.

**Figure 1.**
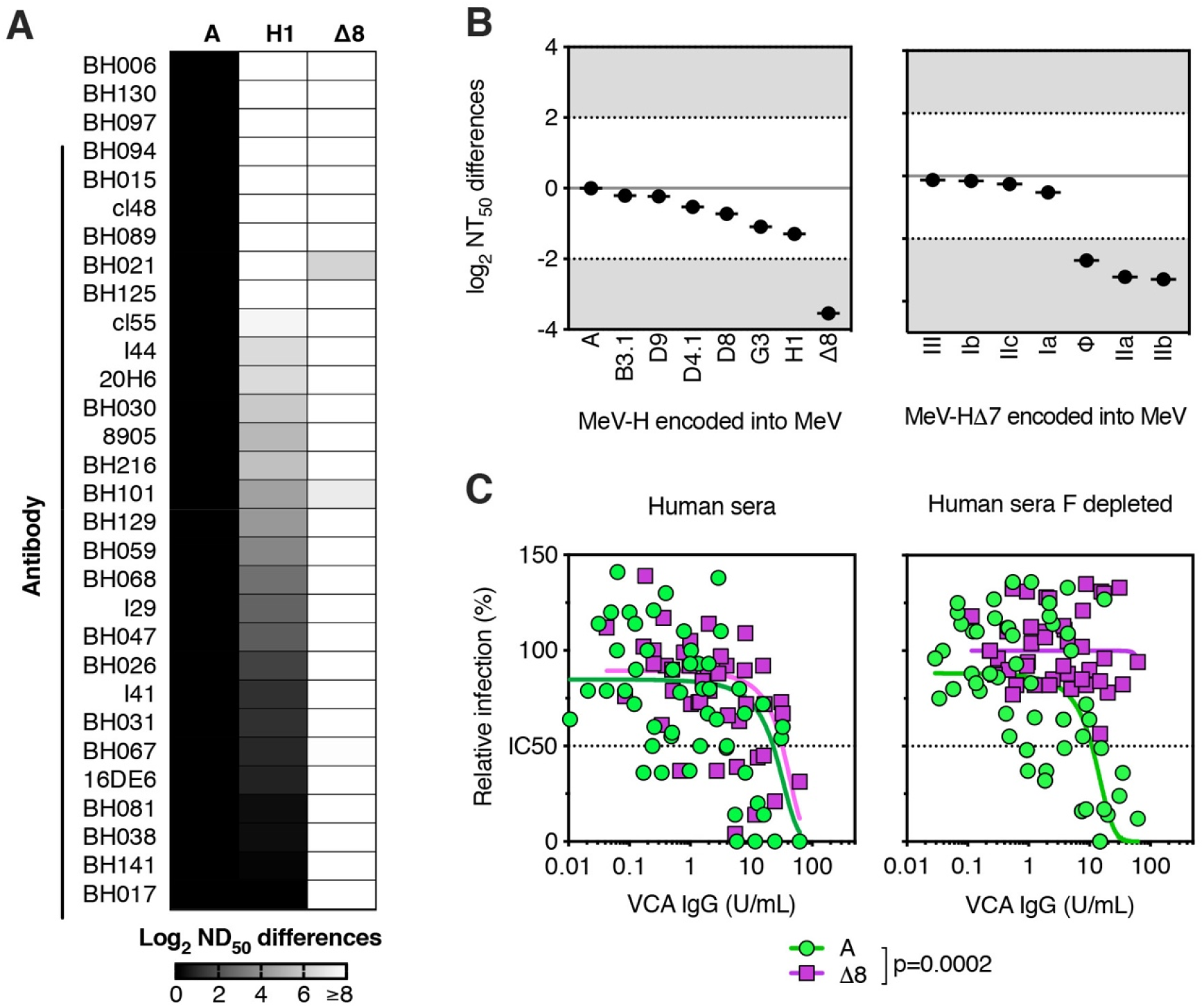
Rational Design of MeV-H Δ8. (A) Neutralization sensitivity of viruses encoding MeV-H A, H1, and Δ8 against a panel of 30 mAbs. Boxes are shaded according to log_2_ ND_50_ reduction. (B, left panel) Neutralization sensitivity of viruses encoding genotype-specific MeV-H (left panel) or (B, right panel) MeV-H Δ 7 mutants (indicated is the antigenic site remaining intact) against mouse sera post-MeV A infection. A difference ≥2 log_2_ (grey-shaded region) is considered antigenically significant. (C) Neutralization sensitivity of MeV A or Δ8 against MeV-immune human sera after depletion of MeV-F-specific antibodies (Fig. S5). Epstein-Barr virus viral capsid antigen (VCA) IgG levels were used to accounting for dilution factors since depletion conditions should not affect their levels.

Interim analysis of the neutralization of mutated viruses by anti-MeV H antisera had revealed that disruption of 4 or fewer antigenic sites did not abrogate polyclonal antibody neutralization (*15–17*). However, in MeV-H Δ8 we observed an 8-fold reduction in the neutralization titer of mouse anti-H antiserum (**Fig. 1B)**. We therefore back-mutated the disrupted antigenic sites to create a panel of Δ7 viruses, each one uniquely presenting a single intact antigenic site. Interestingly, with the exception of the Δ7 viruses retaining antigenic sites Φ, IIa or IIb, all the other four Δ7 viruses were serologically indistinguishable from the parental virus, **(Fig. 1B).** These results were replicated using sera from MeV-H-immunized rabbits (**Fig. S5)**. Thus, simultaneous disruption of at least five antigenic sites is required to manifest resistance to neutralization by MeV-H antisera.

We next sought to determine whether polyclonal sera from measles vaccinees paralleled the above reactivity of mouse and rabbit antisera. Serum samples from Dutch individuals were tested before and after depletion of MeV-F reactive antibodies (**Fig. S6)**. Whereas MeV-F–depleted human serum retained its neutralization potency against viruses displaying MeV-H A, the Δ8 viruses were less readily neutralized (7-fold reduced susceptibility after averaging individual titers) **(Figure 1C).**

Since antigenic site III is known to overlap with the SLAMF1 and NECTIN4 receptor-interacting surfaces of the MeV-H (*18*), the elimination of B cell epitopes in this region was expected to negatively impact binding to these critical pathogenicity determining receptors. We therefore examined the receptor tropisms of the Δ8 variant and of the Δ7 variants exhibiting reduced susceptibility to serum neutralization (**Fig. S7**). None of these viruses was able to mediate infection via SLAMF1, indicating that the emergence of a pathogenic measles strain with reduced susceptibility to serum neutralization would require the virus to reconfigure the receptor interacting residues in antigenic site III in such a way as to destroy the dominant B cell epitopes it shares with existing MV vaccine strains, while retaining NECTIN4 and SLAMF1 receptor binding affinities.

Having demonstrated that the multiply mutated MeV-HΔ8 protein could not interact with pathogenicity-determining MeV receptors, we next sought to eliminate the confounding effect of neutralizing antibodies recognizing the MeV-F glycoprotein (**Fig. 2**). To achieve this, we substituted the MeV-F gene with the corresponding gene from canine distemper virus (CDV). Mutations were introduced into the CDV-F and MeV-H coding sequences to restore and optimize the fusogenicity of this heterologous F/H pairing (**Fig. S8**). This virus, hereafter termed MR (Moraten Resurfaced) was subjected to Sanger sequencing and protein composition analysis to confirm its identity (**Fig. S9**). Propagation of the MeV MR virus on Vero cells was significantly slowed versus the comparable virus incorporating an unmodified MeV-H genotype A protein (**Fig. S10A**). Propagation of MeV-HΔ8 with the parental MeV-F glycoprotein was similarly impaired, indicating that mutation of multiple surface residues in MeV-H compromised protein folding and/or function (**Fig. S10B**).

**Fig. 2.**
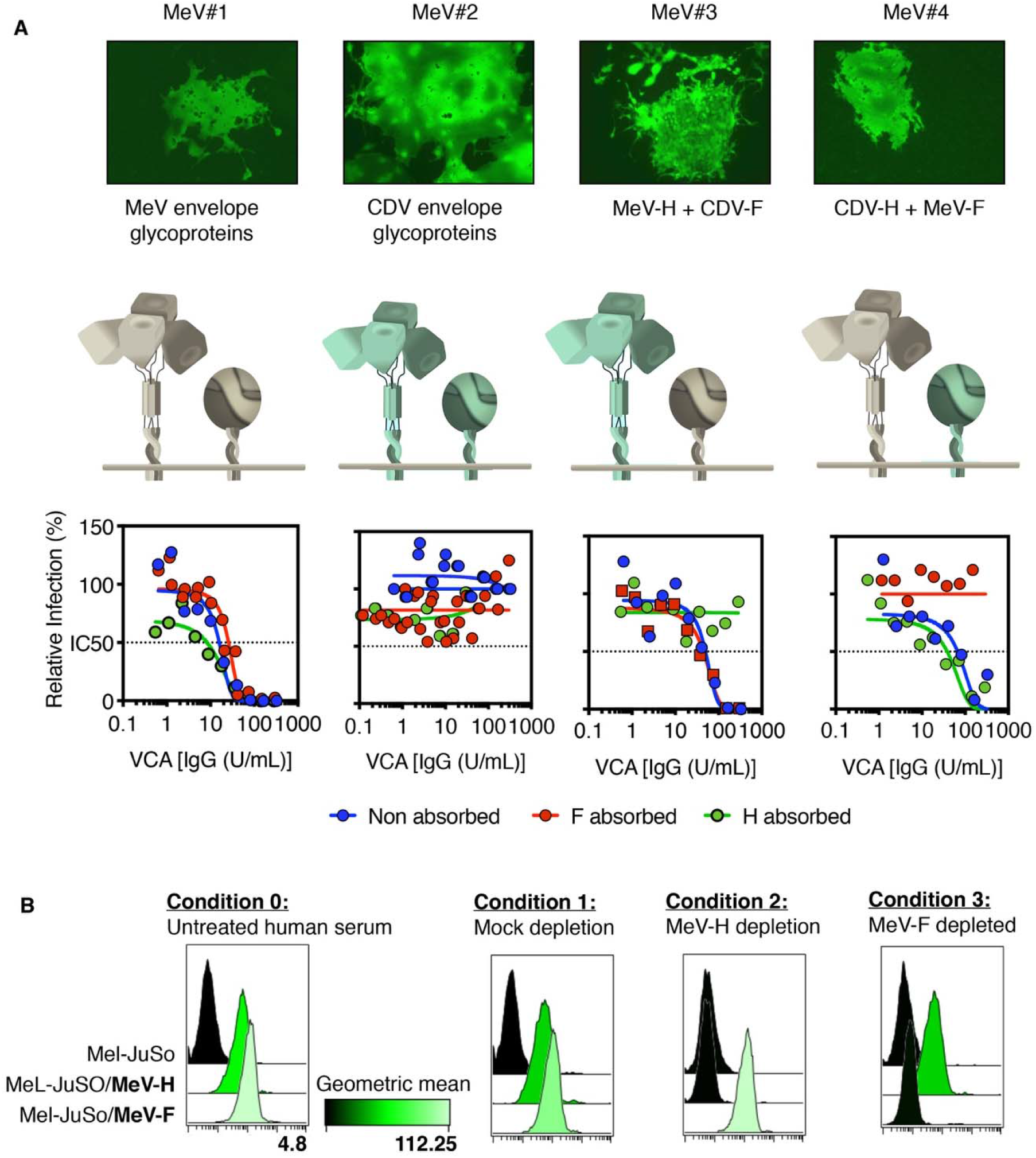
Role of MeV-H and MeV-F in Virus Neutralization. (A) Virus neutralization assay of envelope-exchange viruses. Isogenic recombinant MeV encoding MeV envelope glycoproteins (virus 1), CDV (virus 2), or chimeric viruses (virus 3 and 4) were used to test neutralization sensitivity against pooled human AB sera depleted of antibodies against MeV-H or MeV-F. Representative syncytia are shown. (B) MeV glycoprotein specificity of pooled human AB sera. Conditions and IgG specific levels are described and determined as in Figure S5. The remaining MeV coat-specific antibodies were tested again to confirm successful depletion. Data are shown as histogram plots.

**Fig. 3.**
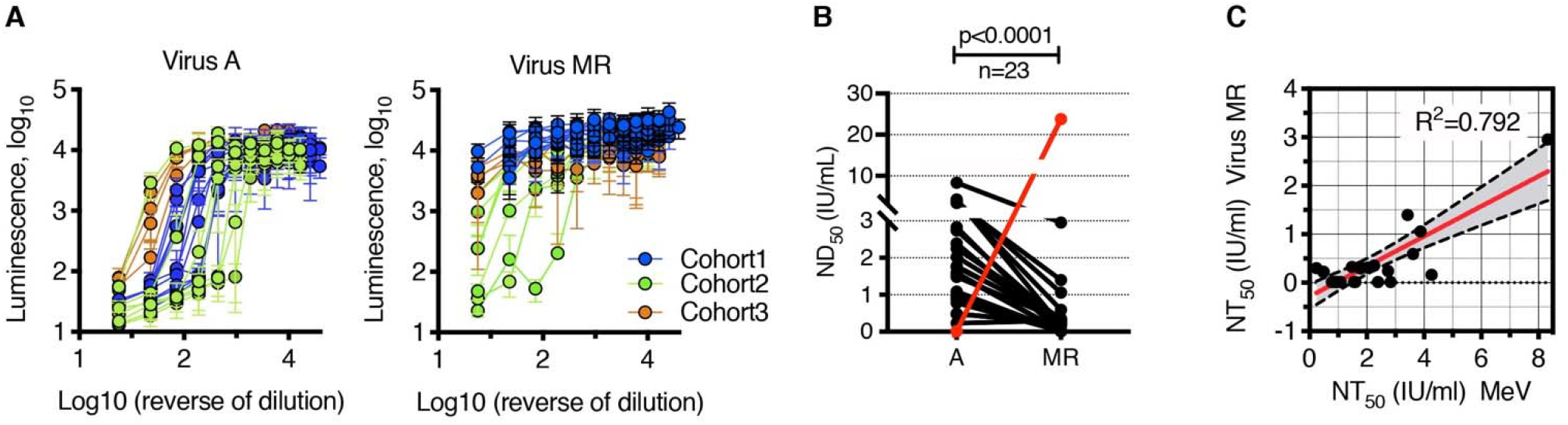
Neutralization activity of human serum samples. (A) Individual neutralization curves. Samples belonging to different cohorts are colored-code. (B) ND_50_ values of MeV-immune human sera against the MeV A and MR. Each line represents an individual sample (N=23). The red line shows ferret serum anti-CDV, used as a control for neutralization. Statistical significance was inferred by a two-tailed paired t-test. (C) Correlation between ND_50_ for the vaccine virus and the MR virus. P<0.001, for both Pearson and Spearman correlation test. The red curve line represents the linear regression line, with dotted lines indicating the 95% CI for the regression analysis.

**Fig 3.**
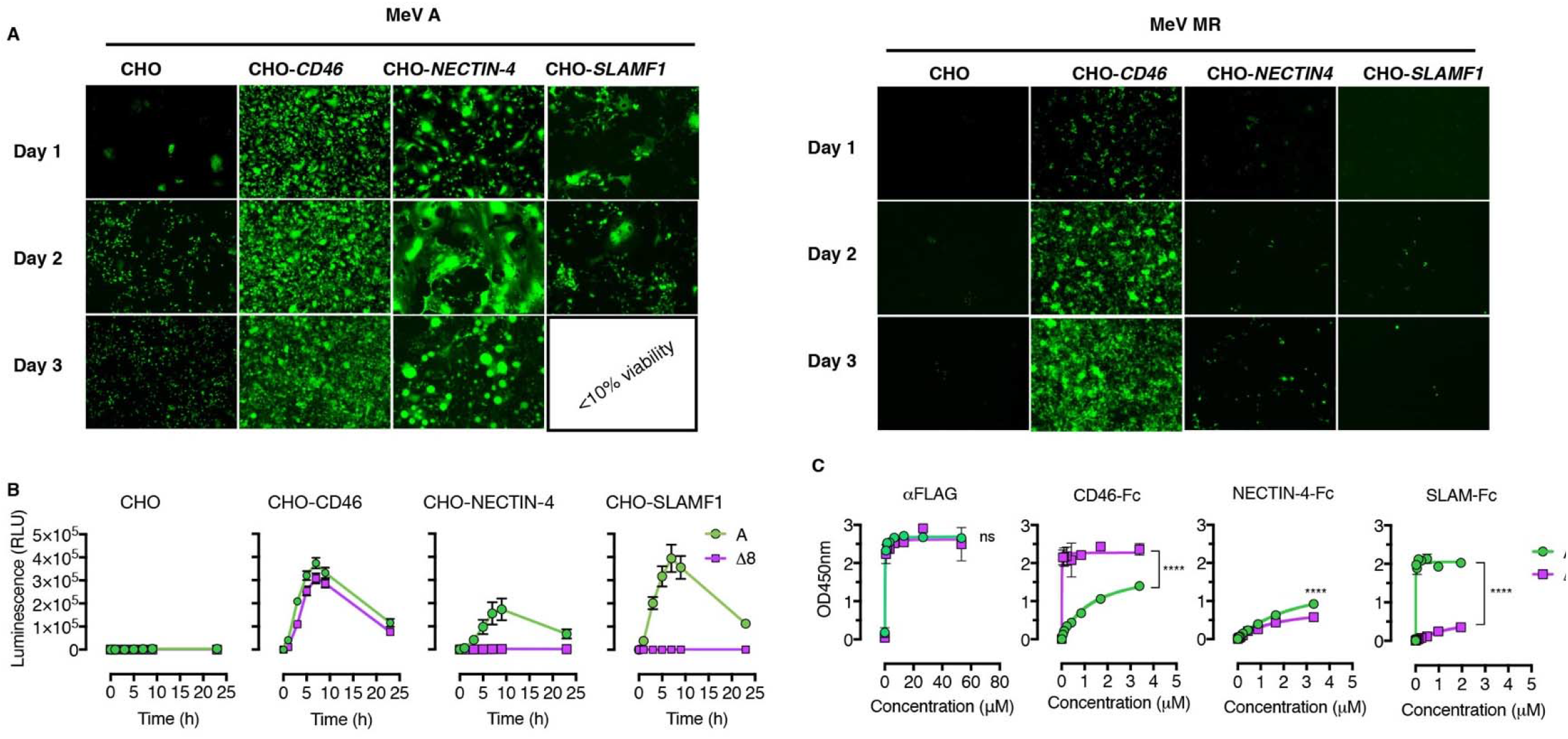
Receptor Usage. (A) CHO cells expressing different MeV receptors were infected at MOI of 1. Images were obtained 3 days after infection. Magnification ×40. (B) Kinetics fusion assay after coexpression of MeV-F with either MeV-H A or Δ8. Mean ± SD. (C) Binding of MeV receptors-Fc to MeV-H protein, monitored by OD. The FLAG epitope present in both MeV-H was used as a coating control. Data are presented as mean ± and were fitted to a 1-site mode of total binding (R 0.99). Statistical significance was determined using the Holm-Sidak multiple comparison test: ns, no significant; **** *P*<.001.

We next tested whether MeV MR was resistant to human serum from vaccinated Dutch (n=13, cohort 1), Minnesotan (n=6, cohort 2) and Hispanic individuals (n=4, cohort 3) using an improved luciferase-based infection neutralization assay (**Fig. S11** and **Fig. 2A**). ND_50_ values of the tested serum samples gave an overall geometric mean neutralization titer 4 log_2_-fold lower MeV-MR versus MeV-A (**Fig. 2B**), suggesting that resistance to neutralization of the MeV MR is fully manifest only at or below a MeV-A ND_50_ titer of 1591 mIU/mL (Fig. 2C).

Interestingly, measles-immune serum does retain some level of neutralizing activity against the MR virus, suggesting that it may also contain protective antibodies directed against subdominant epitopes in the H glycoprotein. To test this, we inoculated the MR or A viruses into immunocompetent HuCD46Ge-IFNar^KO^ mice, harvested sera 4 weeks later and tested for the presence of IgG antibodies directed against the nucleocapsid (MeV-N) or against the MeV-H proteins of pathogenic MeVs (**Fig. S12**). Interestingly, the data confirm that antisera raised against MeV MR do weakly crossreact with wild type MeV-H, indicating that subdominant B cell epitopes may play a significant role in MeV defense. Conversely, antibodies raised against the A virus were able to crossreact weakly with subdominant epitopes in the MeV-H Δ8 protein.

Since MeV-MR is partially resistant to neutralization by measles-immune human sera, it was important to confirm that, like MeV-Δ8, it lacks the ability to use the pathogenicity-determining receptors SLAMF1 and NECTIN4, and enters cells exclusively via CD46. This was confirmed using CHO cells expressing CD46, NECTIN4, or SLAMF1 where, unlike MeV-A, MeV-MR infected only cells expressing CD46 (**Fig. 4A** and **Fig. S13**). This selective tropism is particularly interesting because previous reports claimed that NECTIN4 tropism could not be eliminated independently of CD46 tropism (*19, 20*) (**Fig. S13**). We therefore measured the densities of CD46 and NECTIN4 receptors on our respective CHO cell transfectants and found them to be equivalent (**Fig. S1B**). Cotransfecting plasmids encoding MeV-F and MeV-H Δ8 confirmed that intercellular fusion occurred only in CD46-positive and not NECTIN4-positive CHO cells (**Fig. 4B**) and was similar with CD46 of nonhuman primate origin (**Fig. S15**).

Further mechanistic studies into the discrimination of CD46 over NECTIN-4 showed that MeV-H Δ8 bound more strongly to CD46 than to NECTIN4, and negligibly to SLAMF1. This contrasted with the binding pattern for MeV-H A **(Fig. 4C)** and suggested that MeV-H Δ8 discriminates between CD46 and NECTIN4 via differences in its binding affinities to each of these receptors. We identified no second-site mutations in known contact residues to explain this unexpected segregation of CD46 and NECTIN4 tropisms and therefore postulate that the phenotype may be partially attributable to specific noncontact residues in the MeV-H protein of genotype H1.

Measles-immune human serum is known to negate seroconversion in infants during the first year of life and negates the therapeutic effect of systemically administered oncolytic MeV. Hence, we sought to investigate the impact of passive immunization on the infectivity of the MR virus versus the MeV vaccine. Whereas passive immunization with MeV antisera led to a decrease in luciferase signal from MeV A, there was no reduction in the case of the MeV MR (Fig. S16). Accordingly, systemically administered MeV MR in tumor-bearing mice could reach its target (the tumor cells), in the presence of passive antibodies (Fig. S17).

In summary, by engineering the surface glycoproteins of a MeV vaccine, we have elucidated a number of critical factors determining the remarkable antigenic stability of this monotypic virus. First, MeV has numerous immunologically codominant antigenic sites on its H and F surface glycoproteins. Second, antibodies to each of the seven known antigenic regions on the H glycoprotein are capable of neutralizing virus infectivity. Third, MeVs retaining even a single immunodominant antigenic site on the H glycoprotein remain fully susceptible to neutralization by measles immune human serum. And fourth, the receptor binding surface of the MeV-H glycoprotein is itself an immunodominant antigenic site. Hence MeVs cannot escape their susceptibility to neutralization by measles-immune human serum unless they also lose their tropism for the pathogenicity-determining receptors SLAMF1 and NECTIN4.

Given that a minimum of 5 immunodominant antigenic sites must be disrupted to impact the susceptibility of MeV-H to serum neutralization and that the error rate of the MeV polymerase is approximately 1×10^-5^ mutations per site per round of genome replication (*21*), we estimate the probability of such a virus arising to be of the order of 1 in 10^25^ progeny virions (weighing approximately 10,000,000 kilograms)(*22*). Further, the simultaneous disruption of fewer than five major antigenic sites would confer no selective advantage on the virus, making stepwise evolution an unrealistic pathway to achieve a neutralization-resistant phenotype. However, even a virus insensitive to anti-H antibodies would still be efficiently neutralized by MeV-F-specific antibodies in immune human sera and, even more critical, would lack the SLAMF1 and NECTIN4 receptor tropisms required for pathogenicity and transmission. We therefore conclude there is a near-zero probability for the accidental emergence of a pathogenic MeV capable of evading vaccine-induced immunity.

## Supporting information

Peer Review File

## Acknowledgments

We sincerely thank the following individuals: Mark J. Federspiel, PhD, for mAb cl48. Ianko D. Iankov, MD, PhD, for mAb 20H6 and Professor Claude Muller, MD, for the remaining murine nAb as well as discussions. Roberto Cattaneo, PhD, for Vero/dogSLAM, measles virus antigenome plasmids and rabbit anti-MeV antibodies for Western blotting. Rik L. de Swart, PhD, for the Mel-JuSo cell lines, human serum samples and antibodies C28-10-8 and F3-5. Mayo Clinic Biobank, for serum samples from the Minnesotan cohort. M. Cristine Charlesworth, PhD, and Benjamin J. Madden (Mayo Clinic Proteomics Core) for purification and mass spectrometry of soluble receptors-Fc. Zene Matzuda, MD, PhD, DSc, for the split luciferase plasmids. Eugene Bah, for his assistance with the initial quantitative fusion assays and useful discussion. We also thank the Mayo Clinic Biosafety Committee for their critical reading of the manuscript and helpful discussion.

## Author contributions

Conceptualization: M.A.M.-A., S.J.R.; Methodology: M.A.M.-A., S.J.R.; Validation: M.A.M.-A. S.J.R; Investigation: M.A.M.-A., and LZ.; Analysis: M.A.M.-A., S.J.R.; Resources: M.A.M.-A., S.J.R.; Writing, review and editing: M.A.M.-A., S.J.R. Funding acquisition: S.J.R. Approval of the final manuscript: All authors.

## Data and materials availability

All data needed to evaluate the conclusion of this paper are present in either the main text or the supplementary materials.

## Supplementary Materials

Materials and Methods

Figures S1-S17

