## Supplementary Information for "Pathogenic measles viruses cannot evolve to bypass vaccine-induced neutralizing antibodies"

Although the modified envelope glycoprotein complex generated in this project could be theoretically implanted into a pathogenic measles virus, doing so would destroy its ability to interact with SLAMF1 and NECTIN4, the two receptors used by pathogenic measles strains (and by all other pathogenic morbilliviruses) to cause disease. To restore these pathogenicity-determining tropisms, a major antigenic site would be reintroduced into the H glycoprotein of the the hypothetical virus, thereby rendering it virus fully susceptible to measles-immune human serum.

**Materials and Methods**

Prior to the onset of experimentation, the Mayo Clinic Institutional Biosafety Committee (IBC) performed a Dual Use Research of Concern (DURC) evaluation on the proposed project. Although it flagged as a project with DURC elements, the project was allowed to continue with work occurring in a certified Biosafety Level 2/2+ facility and with ongoing oversight by the Biosafety Office.  Despite tremendous mutational pressure, evidence that there are mutations in the viral H and N genes, and widespread infections throughout geography and time, the virus has remained monotypic. Given those elements, it was the view of the authors and the institutional review team that there was an unknown factor protecting the monotypic status of the vaccine strain which decreased the concern that the work proposed by this project would result in a virus that could pose increased public risk. Had evidence arising during the course of the project that would have altered that determination, an additional review of the project would have been conducted.

*Cells and Viruses*

Vero cells (CCL-81, ATCC) and Vero cells stably transfected with human

SLAMF1 (*1*) or with its canine counterpart (*2*) were grown in Dulbecco’s modified minimal essential medium (DMEM) (HyClone, GE Healthcare Life Science) supplemented with 5% (vol./vol.) heat-inactivated fetal bovine serum (Gibco) and 0.5 mg/mL of Geneticin (G418; Corning) for Vero/hSLAM or 1 mg/mL Zeocin (ThermoFisher) for Vero/cSLAM. CHO cells, CHO-CD46 (*3*), CHO-SLAM (*4*) and CHO-NECTIN4 (*5*) were cultured as previously described. Baby hamster kidney cells were maintained in DMEM-10% fetal bovine serum. The human melanoma Mel-JuSo cell line and derivatives, transfected with the full-length MeV-F and MeV-H genes (Mel-JuSo-F or Mel-JuSo-H) were cultured in RPMI 1640 medium (Corning, Manassas, VA) supplemented with G418.

Viruses have been named according to the genotype of MeV-H gene expressed and were propagated as described previously (*6, 7*). For mouse inoculation, viruses were purified through a sucrose gradient as described elsewhere (*8*).

*Constructs and Rescue of Recombinant MeVs*

Viruses used in this study were derived from the molecular cDNA clone of the Moraten/Schwartz vaccine strain. For the construction of recombinant MeVs, plasmid pB(+)MeV^vac2^(eGFP)N plasmid (*9*), encoding for a rMeV expressing the enhanced green fluorescent protein (eGFP) upstream of the N gene, and plasmid pB(+)MeV^vac2^(ATU)P, plasmid encoding for a rMeV with an additional transcription using downstream of the phosphoprotein gene (*9*), were used. A DNA fragment encoding for firefly luciferase (GenBank accession #KF926476) with Mlu/AatII restriction sites at its extremities, was then cloned into the corresponding sites of the ATU to obtain pB(+)MeV^vac2^(Fluc)P. To avoid plasmid instability during propagation in bacteria, *Escherichia coli* Stbl2 cells (Invitrogen, 10268019) were used (grown at 30°C).

We used the CDV Ondersterpoort vaccine strain H (CDV-H) and F (CDV-F) genes, originally contained in pCG plasmid (*10*), to produce envelope-exchange MeVs. To replace MeV-H from the MeV backbone, site-directed mutagenesis (QuikChange site-directed mutagenesis kit, Agilent) was used to remove a SpeI site in CDV-H, and a Y537D substitution was then introduced, which reduces binding by crossreactive nAb in human sera (*11*). PacI and SpeI restriction sites (underlined) were introduced into the beginning and end of the gene, respectively, by polymerase chain reaction using forward primer 5′‑ttaattaaaacttagggtgcaagatcatcgataatgctcccctaccaagacaagg‑3′ and reverse primer 5′‑actagtgggtatgcctgatgtctgggtgacatcatgtgattggttcactagcagccttaatggtggtgatggtggtggctcccccttgcggccgcggccggctgggccgctctaccctcgatacggttacatgagaatcttatacggac‑3′, leaving the untranslated region unchanged. The PCR product was digested with PacI and SpeI and cloned into the MeV backbone. To replace MeV-F from the MeV antigenome plasmid, pCG-CDV-F was digested with HpaI/SpeI and inserted into similarly digested pCG-MeV-F. The NarI/SpeI fragment of this plasmid was then used to replace that of MeV.

The recovery of recombinant MeVs was performed as previously described (*12, 13*).

*Fusion Assay*

Cells (5×10^5^ cells/well in a 6-well plate) were cotransfected using Fugene HD (Promega) with 1 μg of pCG plasmid encoding vaccine strain MeV-F and pCG encoding the appropriate MeV-H. Fusion activity was evaluated 24 hours later with Hema-Quik staining (Fisher Scientific #23123745).

To quantify cell fusion, we used the dual-split luciferase assay as previously described (*14*). Briefly, effector baby hamster kidney cells (3×10^4^) in black 96-well plates were cotransfected with 33 ng each of the MeV-H and MeV-F expression plasmids and one of the split luciferase plasmids, DSP_8-11_. As a control, only the MeV-F and DSP_8-11_ plasmids were transfected. CHO cells and CHO cells expressing the respective MeV receptors (2×10^5^ cells/well in 6-well plates) were transfected with 1.5 μg of the other dual-split-reporter plasmid (DSP_1-7_). Twenty-four hours after transfection, target cells were detached with Versene (Life Technologies) and cocultured with the effector cells in Fusion media (DMEM-F12 without Phenol Red + 40 mM HEPES), supplemented with 1:1000 dilution of the cell-permeable luciferase substrate EnduREN (Promega). Luminescence resulting from cell fusion and mixing of cytoplasmic content between target and effector cells was monitored with a Topcount NXT Luminometer (Packard Instrument Company, Meriden, CT) at the indicated time points. The data shown are the mean and standard deviation of 3 replicates for each H plasmid.

*Fluorescence-Activated Cell Sorting Analysis and Quantification of Cell Surface Molecules*

Cells were washed and detached by using Versene (Gibco) and immediately incubated with phycoerythrin-conjugated antibodies: anti-SLAMF1 (FAB1642P; R&D Systems), anti-CD46 (FAB2005P; R&D Systems), anti-NECTIN4 (FAB2659P; R&D Systems), or control isotype antibody (IC0041P; R&D Systems). After a 1-hour incubation at 4°C, cells were washed again and fluorescence was measured in a FACSCanto flow cytometry system (BD Bioscience). The number of receptors per cell was estimated with calibration beads (BD QuantiBRITE; BD Biosciences) as the reference standard.

*Recombinant Proteins and Binding Assays*

The coding sequence of the CD46 ectodomain (residues 35-328) was amplified via PCR from pGEM-CD46 vector (Sino Biologicals Inc., HG12239-G) and inserted into a pFUSE vector (pfc1-hg1e3; Invivogen) in frame with the murine Ig κ-chain leader sequence and a 3C protease cleavage sequence at the 5′ end of the Fc region using In-Fusion cloning kit (Clontech). CD46-Fc, SLAM-Fc, and NECTIN4-Fc (*7*) recombinant proteins were expressed in Expi293F cells (ThermoFisher) and purified from culture supernatant as previously described (*7*).

The expression and purification of recombinant soluble MeV-H were performed as previously described (*15*).

Binding of the receptors-Fc to MeV-H was determined by enzyme-linked immunosorbent assay as previously described (*7*). The absorbance at 450 nm was measured with an Infinite M200Pro microplate reader (Tecan). The data were analyzed using Prism software (GraphPad) and adjusted to a 1-site binding saturation mode to determine the half-saturating concentration (apparent Kd values [dissociation constant]). Values reported showed excellent fit (R^2^>0.99).

Recombinant nucleocapsid protein was purchased from CD Creative Diagnostics (NY, USA).

*Virus Protein Content and Cell Surface expression of MeV-H proteins*

Virus preparations were heated in the presence of dithiothreitol, fractionated into 4-12% Bis-Tris polyacrylamide gel, and transferred to polyvinylidene fluoride membranes. Blots were analyzed with anti-MeV-Hcyt, anti-MeV-N, anti-MeV-F, anti-GFP antibodies and mouse anti–β

-actin (*7, 16*) and probed with a conjugated secondary rabbit antibody (ThermoFisher, #31642). The blots were incubated with SuperSignal West Pico chemiluminescent substrate (ThermoFisher) and analyzed with a ChemiDoc Imaging Sytem (Bio-Rad). For cell surface expression of MeV-H proteins, MelJuSO cells were transfected with 2.5 μg of the corresponding MeV-H expression plasmid, and cells were labelled 24 hours later with EZ Link^TM^ Sulfo-NHSS-SS-Biotin, followed by quenching and lysis of the cells according to the manufacturer’s instructions (ThermoFisher, Cat# A44390). Biotinylated proteins were immunoprecipiated with NeutrAvidin and eluted by boiling in SDS-PAGE buffer containing dithithreitol (DTT). Protein were next fractionated and blotted as indicated above with anti-MeV-Hcyt and anti–β

-actin antibody.

*Passive Immunization and In Vivo Imaging*

Neutralizing antibodies (600 mIU) were administered in the form anti-serum and they were obtained through the NIH Biodefense and Emerging Infections Research Resources Repository, NIAID, NIH: Polyclonal Anti-Measles Virus, Edmonston, (antiserum, Guinea pig), NR-4024; and polyclonal anti–CDV, Lederle avirulent (antiserum, ferret), NR-4025. We have previously determined their specificity and potency of the reagents (*16*). HuCD46Ge-IFNAR^KO^ mice (*17*) received intra-peritoneally virus anti-serum 3 hours before injection of 2x10^5^ TCID_50_/100μl of rMeV(FlucP) viruses, following the same route. At 24, 48, 72 and 96 hours, mice were anesthetized and inoculated with D-Luciferin (GoldBio Cat.# LUCK-100) 15 min before *in vivo* imaging of bioluminescence using an IVIS Spectrum instrument (Perking Elmer). Female CB17 ICR SCID mice of 4-6 weeks old were purchased from Taconic (Germantown, NY) and implanted subcutaneously with 1x10^7^ KAS 6/1 cells one day after whole body irradiation (2Gy). When tumor reached a volumen of 0.5 cm^3^, 2x10^5^ TCID_50_ units of rMeV(FlucP) viruses or 100μl of PBS were injected through the tail vein. *In vivo* imaging was then recorded as described previously.

*Serologic Assays*

Virus neutralization assays were based on the fluorescence-based PRMN assay, as previously described (*6, 7, 15*). Alternatively, a luciferase-based neutralization assays was developed. After incubation for 72 h at 37°C and 5% CO_2_, 100μl of DMEM/F-12 (ThermoFisher Scientific, Cat.#11330-021) containing 0.5mM of D-Luciferin was added to each well of a black 96-well plate. Luciferase-expressing cells were quantified with an Infinite M200 Pro multimode microplate reader (Tecan Trading AG). Each assay was repeated at least twice and on different days, with 4 replicates per assay. Fifty percent inhibitory concentration (IC_50_) was calculated after fitting the data to a sigmoidal dose-response (variable slope) with GraphPad software (Prism 7). Values were converted to mIU/mL by using the third World Health Organization international antimeasles standard, as previously described (*18*).

Rabbit anti-MeV-H sera were generated by immunization with adenovirus expressing the MeV-H from the vaccine strain (*19*).

Murine monoclonal anti-MeV-H antibodies were produced and characterized as previously described (*6, 20-24*). Polyclonal antibodies were generated by gene-based hydrodynamic injection (*25*) of C57BL/6 mice with 20μg of plasmid DNA or by intraperitoneal inoculation of HuCD46Ge-IFNAR^KO^ mice with 1 dose of 10^5^ FFU of MeV-expressing GFP. Serum was obtained 4 weeks after inoculation.

Human serum samples were from 3 cohorts. Cohort 1 samples were obtained from the Erasmus Medical Center serum bank and have been previously described (*26*). Members of cohort 1 likely were never exposed to wild-type MeV and likely received a monovalent measles vaccination at age 14 months and a measles-mumps-rubella (MMR) vaccination at age 9 years. Cohort 2 samples were obtained from the Mayo Clinic Biobank in Olmsted Country (Rochester, MN); this region has no documented evidence of circulating measles or rubella virus in the past and has high vaccine coverage. However, MMR vaccination status was not available for most samples. The subjects’ age for cohort 2 ranged from 30 to 44 years, and 80% were female. Serum samples were tested by enzyme-linked immunosorbent assay for IgG antibodies specific to MeV and rubella virus, and no differences were identified between those with 1 documented dose of MMR and those without vaccination documented. Cohort 3 samples were custom purchased from Innovative Research Inc, based on age and race. Hispanic donors between 20 and 29 years old were chosen to increase the likehood of obtaining serum samples from vaccinated people. The Latin America region was the first in the world to have eliminated measles and people in that age range were considered to have acquired their MeV protection exclusively through vaccination without further boosting of immunity through exposure to wild-type viruses. All serum samples were heat inactivated.

The Epstein-Barr virus IgG titer was determined by a commercially available assay (IBL International GMbH, cat. No. 57351). The assay for determining MeV-specific IgG levels was described previously (*26, 27*). Estimation of MeV-N antibodies were performed by enzyme-linked immunosorbent assay using a nucleocapsid protein as the antigen. A standard calibration curve was generated with anti-MeV-N (Millipore; clone 83KKII).

*Structural Modeling*

A model of the MeV-H was generated from the crystallographic structures of the MeV-H alone (PDB 2ZB5) and the MeV-H-SLAMF1 complex (PDB 3ALZ). The model was then

submitted for in silico glycosylation using the GlyPro server (http://www.glycosciences.de), which produced a complex penta-antennary N-glycan model at all predicted N-glycosylation sites, including N168 and N187, that are part of disordered regions. The structures were superimposed and manipulated with PyMOL software (<http://pymol.org>).

*Statistical Analysis*

Statistical significance was calculated with GraphPad Prism 7 following the appropriate statistical test, as described in the text corresponding to each figure.

**Table 1.** Mutations Engineered Into Measles Virus Hemagglutinin to Eliminate All Known B-Cell Epitopes

| **Antigenic Site** | **Alternative Name** | **Mutation** | | |
| --- | --- | --- | --- | --- |
|  |  | **Amino Acid Substitution** | **Monoclonal Antibody Resistance** | **Reference** |
| Φ (*7*) | E1 (*19*) | N282K | BH015, BH130 | (*7*) |
| Ia (*28*) | I (*29*), VI (*30*), E4 (*19*), LE (*31*),  V (*30*) | E235G | E185 | (*30*) |
|  |  | G302R | E39 | (*30*) |
|  |  | Y310C | BH038, BH141, I-29 | (*19*) |
|  |  | Q311R | E103 | (*30*) |
| Ib (*28*) | NE, IV (*30*) | L249P | BH047, BH059, BH129 | (*15*) |
| IIa (*32*) | II (*28, 29*), SSE (*31*), VII (*30*) | E488K | BH097 | (*33*) |
|  |  | G491D | 16CD11 | (*22*) |
| IIb (*32*) | II (*28, 29*), SSE (*31*), VII (*30*) | 416DLS🡪 NLS | BH125,  E128 | (*7*)  (*30*) |
| III (*29*) | IIIA (*28*), IIIB (*28*)  VII (*34*), RBE (*31*), E2 (*19*) | S189P | I-44 | (*22*) |
|  |  | D505T | 80-II-B2 | (*28*) |
|  |  | R533G | Cl55  16DE6 | (*32*)  (*22*) |
|  |  | R547G | 20H6 | (*33*) |
|  |  | F552V | I-41 | (*22*) |
|  |  | R377Q, M378K | L77 | (*35*) |
| ‘Noose’ (*36*) | IV (*28, 29*), HNE (*36*), I (*30*), E3 (*19*), D, D/E (*35*) | P397L | BH006, BH216 | (*37*) |
|  |  | N405S | 8905 | (*32*) |
| IIc | No applicable | E471K | BH030 | Current study |

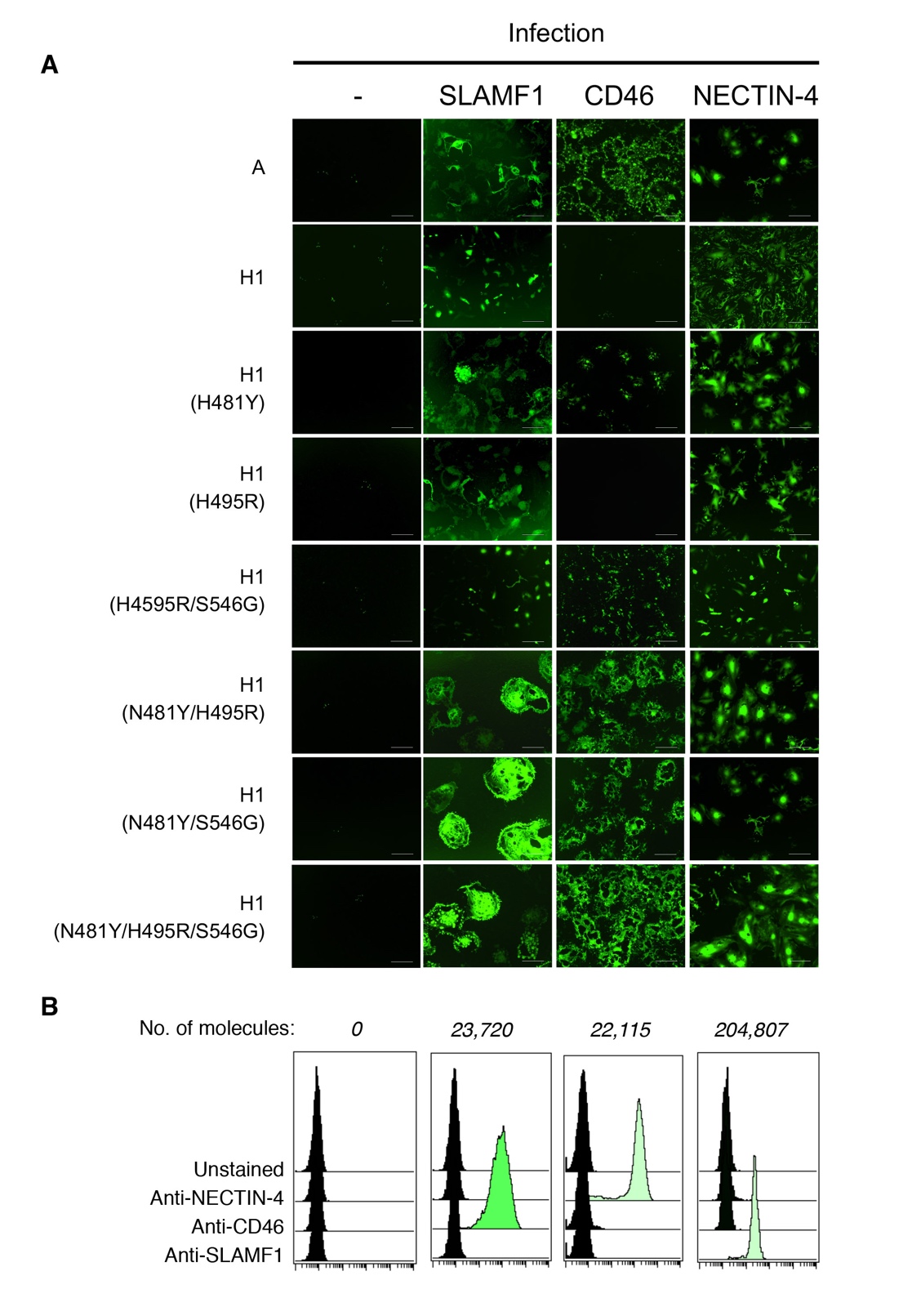

**Sup. Fig. 1.** Engineering CD46 tropism into a wild-type MeV-H protein (genotype H1). (A) CHO cells expressing cellular receptors SLAMF1, CD46, or NECTIN4 and control cells were infected with eGFP-expressing MeV at MOI 0.1. eGFP autofluorescence was recorded 72 hours after infection. Mutations and the corresponding genotype background are indicated. (B) Quantification of cell surface expression for the receptors. Cells were stained with PE-conjugated antibodies and the number of molecules per cell was determined with QuantiBRITE BD.

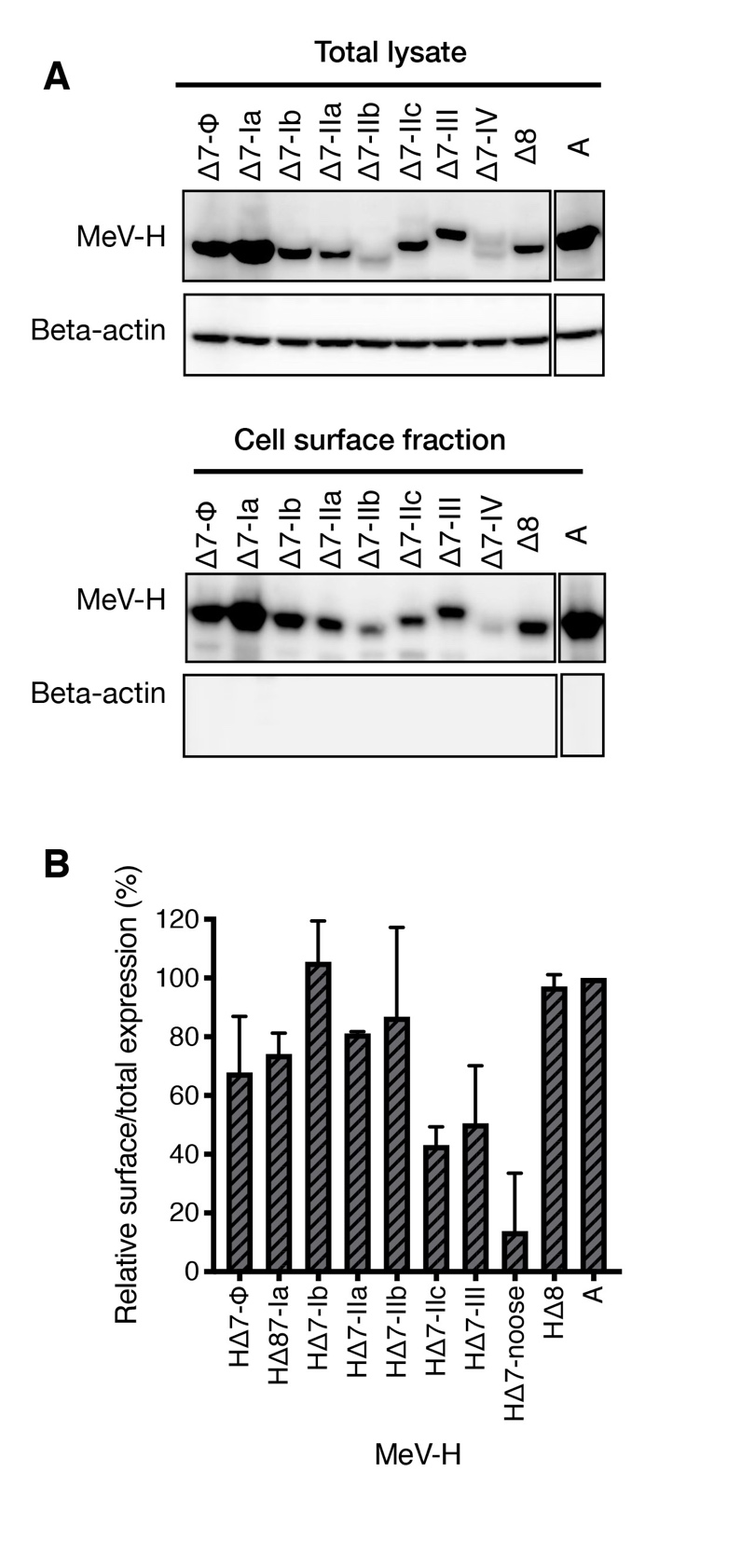

**Fig. S2.** Relative surface expression of MeV-H mutants. Cells transfected with the indicated MeV-H expression plasmid were biotinylated 24 hours later according to manufacturer’s instruction and lysed with RIPA buffer. Clarified supernantants were immunoprecipited (IP) with NeutrAvidin agarose and fractionated by SDS-PAGE into a NUPAGE 4-12% Bis-Tris Gel. Both IP fraction (cell surface expression) and whole cell lysate (total expression) were probed with anti-MeV-H cyt or β -actin antibodies. Densitometry of the corresponding bands were determined from two independent gels and plotted as the amount of MeV-H in the surface versus total. The reader is referred to Fig. S2, panel D for spatial location of the antigenic sites.

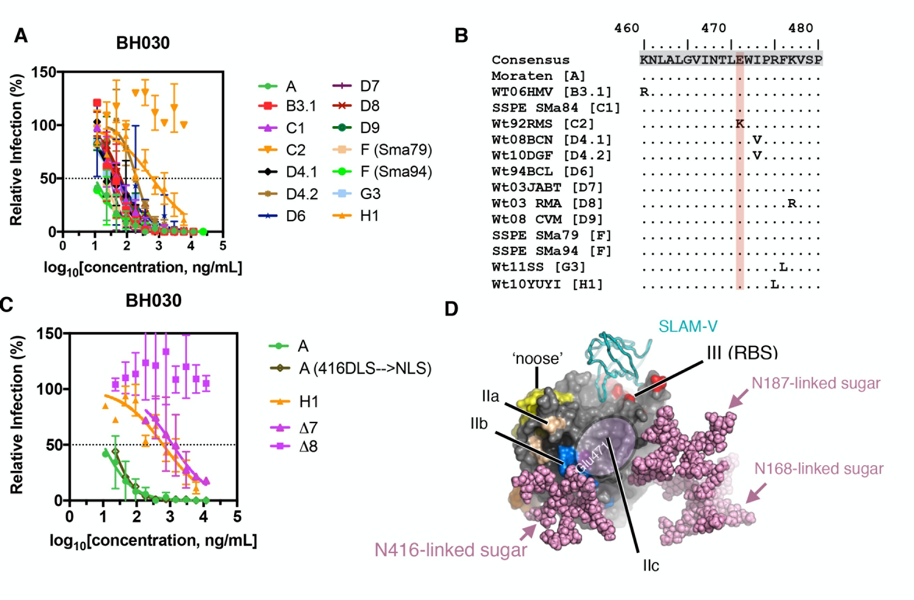

**Fig. S3**. Epitope defined by mAb BH030 corresponds to an operationally distinct antigenic site. PRMN assay of mAb BH030 against recombinant MeV encoding different genotype-specific MeV-H. Of note, C2 viruses selectively escape neutralization by this mAbs. (B) Amino acid sequence alignment for the putative BH030 epitope, showing the differential amino acid substitution in C2 viruses. (C) H1 viruses show resistance against BH030 neutralization in comparison with A viruses and A mutant (416DLS🡪NLS), but are still neutralized, as Δ7 viruses are. The addition of E471K into Δ7, name herein Δ8, results in BH030 escape. (D) Putative antigenic site defined by mAb BH030. Glu471 is not masked by N416-linked sugar, which shields site IIb, and therefore define a putative new antigenic site, IIc, that locates between sites IIb to site III.

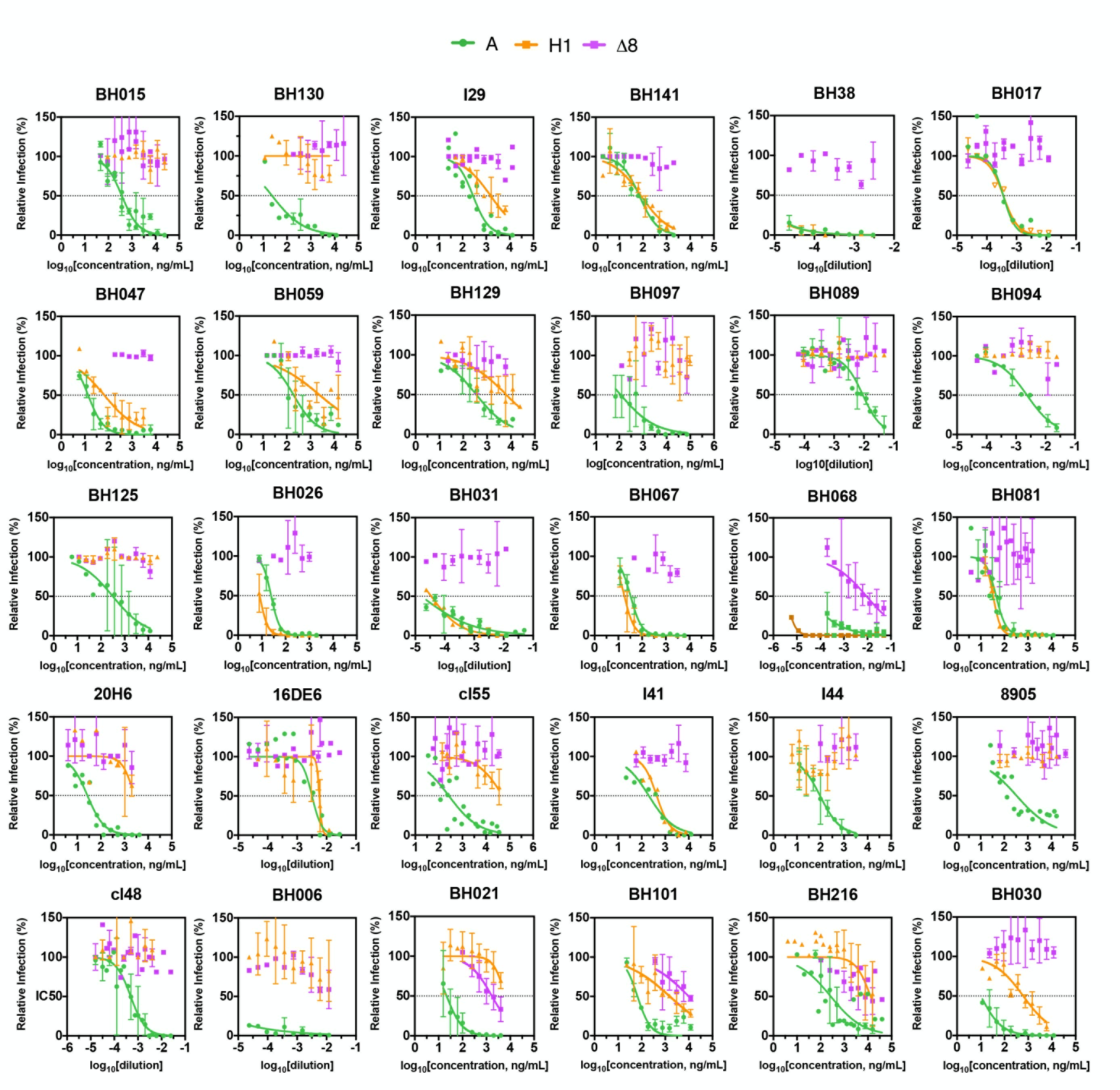

**Fig. S4.** Individual neutralization curves for the panel of 30 murine mAb. PRMN assay of murine monoclonal antibodies was performed against recombinant MeVs encoding MeV-H genotype A, H1 or MeV-H Δ8. After incubation with the indicated mAb, the mix was added to a monolayer of Vero/hSLAM cells. Plaques in the different dilutions were counted and plotted as percentage of infection in the absence of antibody. Data points represent mean ± standard deviation of at least two independent experiments performed in quadruplicate. Data were fitted by non-linear regression analysis with Graph Pad Prism. Inhibitory concentration 50% (IC_50_) is indicated by a dotted line.

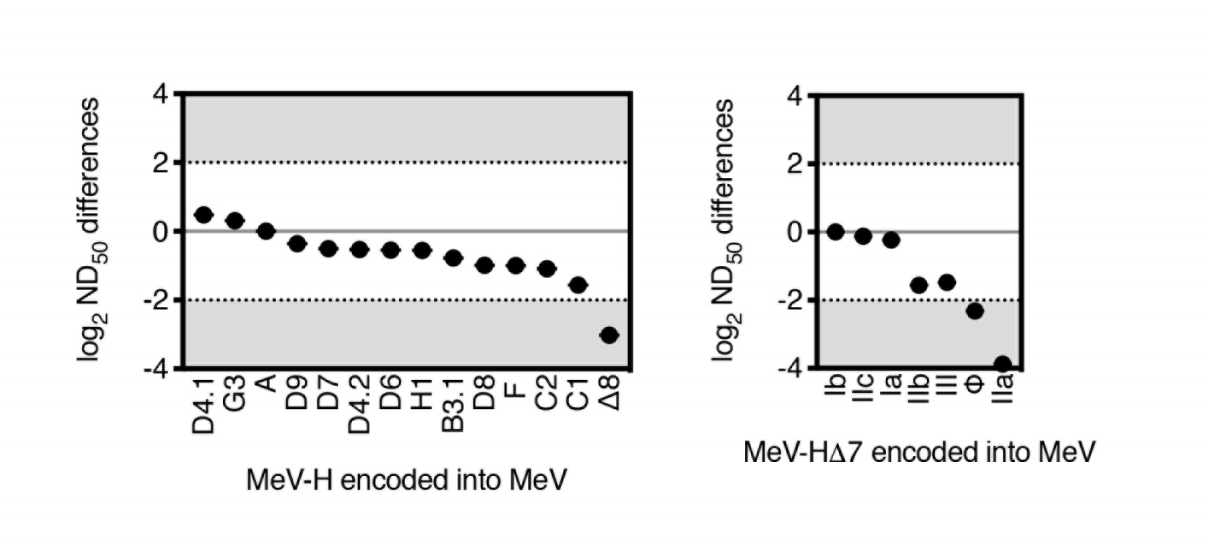

**Fig. S5.** Neutralization sensitivity of viruses encoding genotype-specific MeV-H (left panel) or (right panel) MeV-H Δ 7 mutants (indicated is the antigenic site remaining intact) against rabbit post MeV-H A immunization. The results are reported as in Fig. 1.

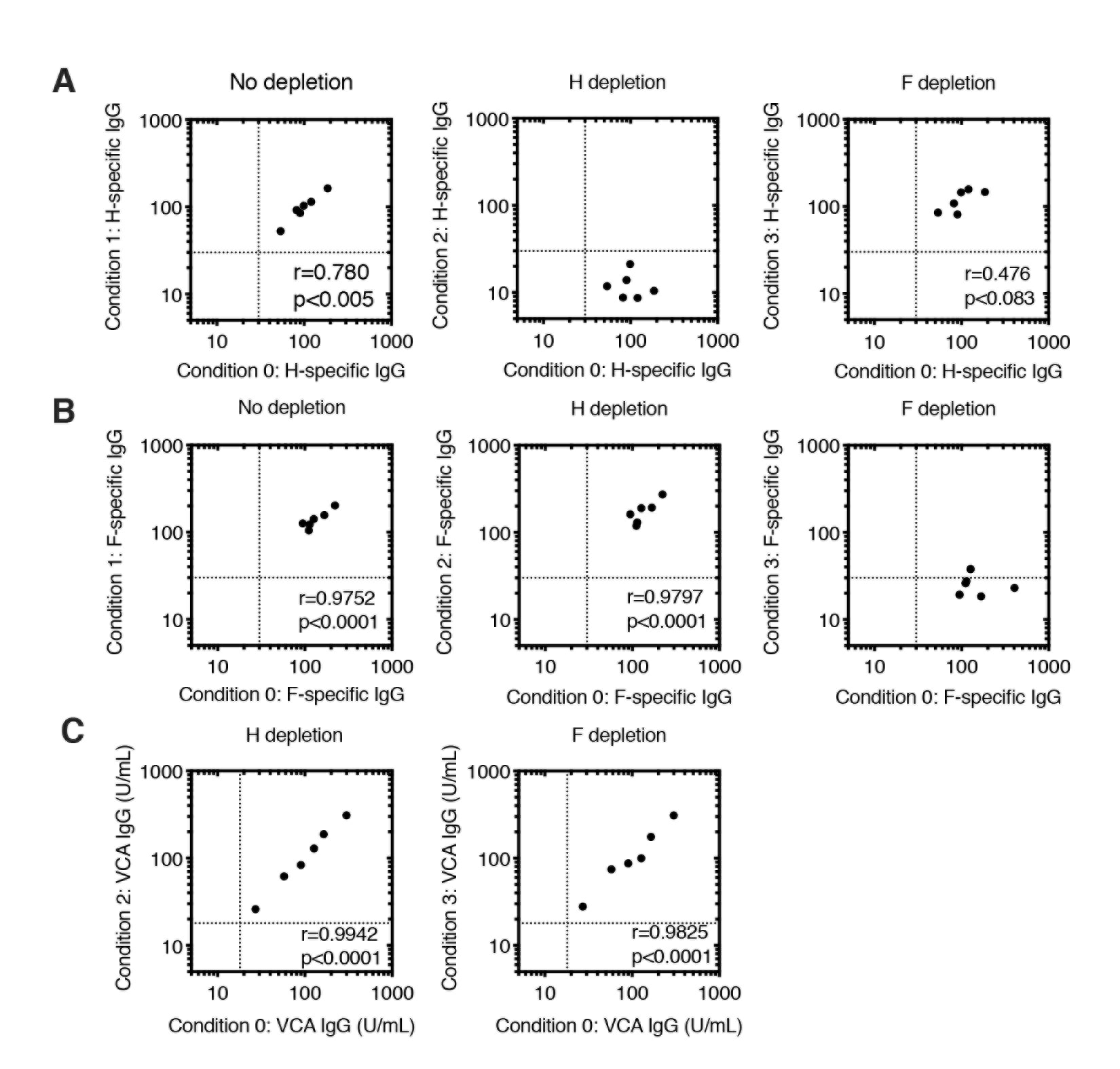

**Fig. S6.** of MeV glycoprotein-specific antibodies. IgG antibody levels after incubating MeV-immune human sera with cells non-expressing (condition 1) or expressing MeV glycoproteins MeV-H (condition 2) and MeV-F (condition 3) and comparing their levels to those of untreated human sera (condition 0). Serum samples were diluted 1:10 in culture medium, applied to a monolayer of Mel-JuSo-F cells, and cultured for 4 days. Supernatants were collected and tested for the presence of specific antibodies against MeV-H (A) and MeV-F (B) by an immunofluorescence assay (fluorescence-activated cell sorting). Values <30 were considered negative for antibody binding (lower crosshairs). (C) IgG levels for the Epstein-Barr virus viral capsid antigen (VCA)

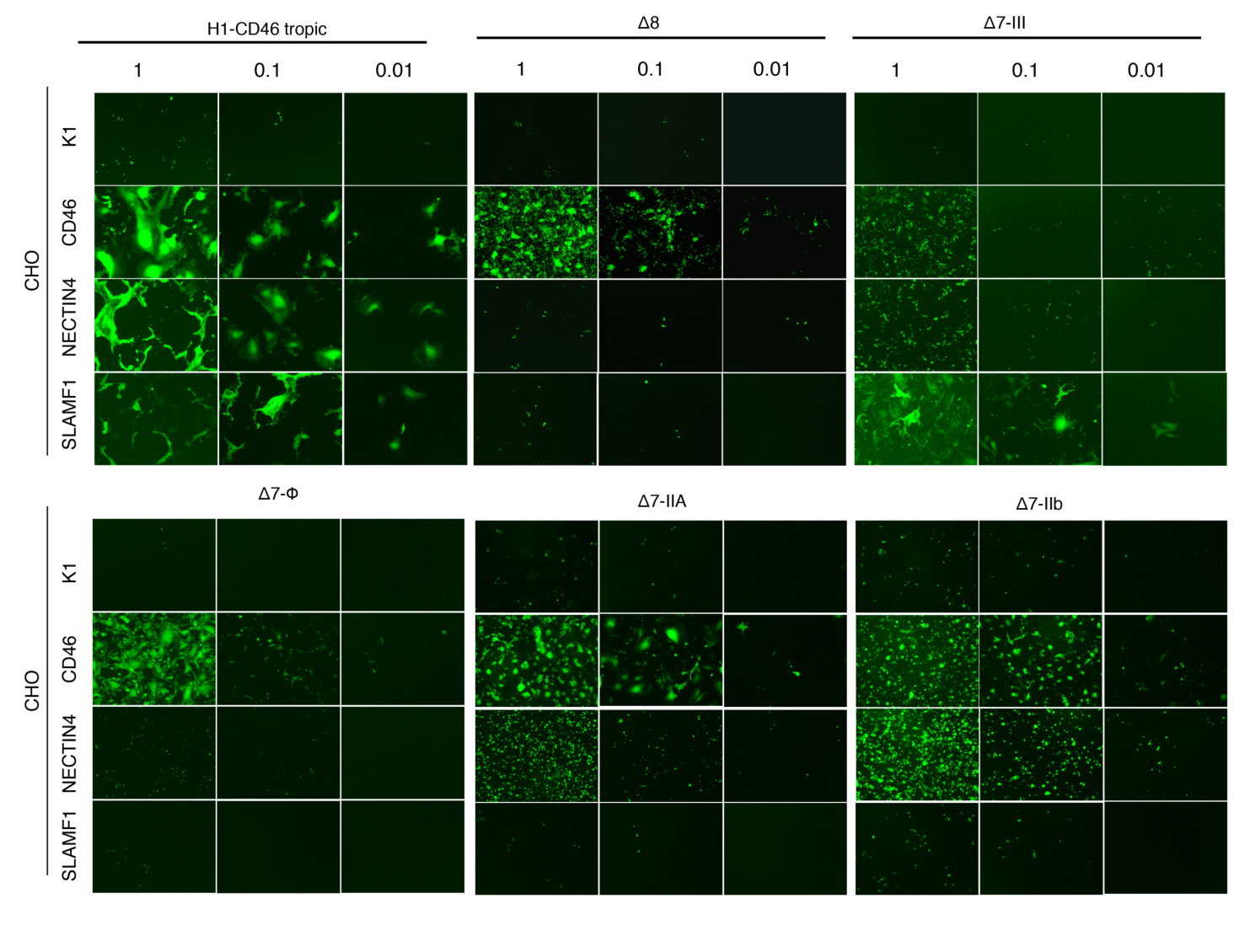
**Fig. S7.** Tropism of recombinant MeV encoding parental or mutated MeV-H. CHO cells expressing the different MeV receptors were infected at the indicated MOI. 48 hours later, EGFP autofluorescence was recorded. Magnification x40.

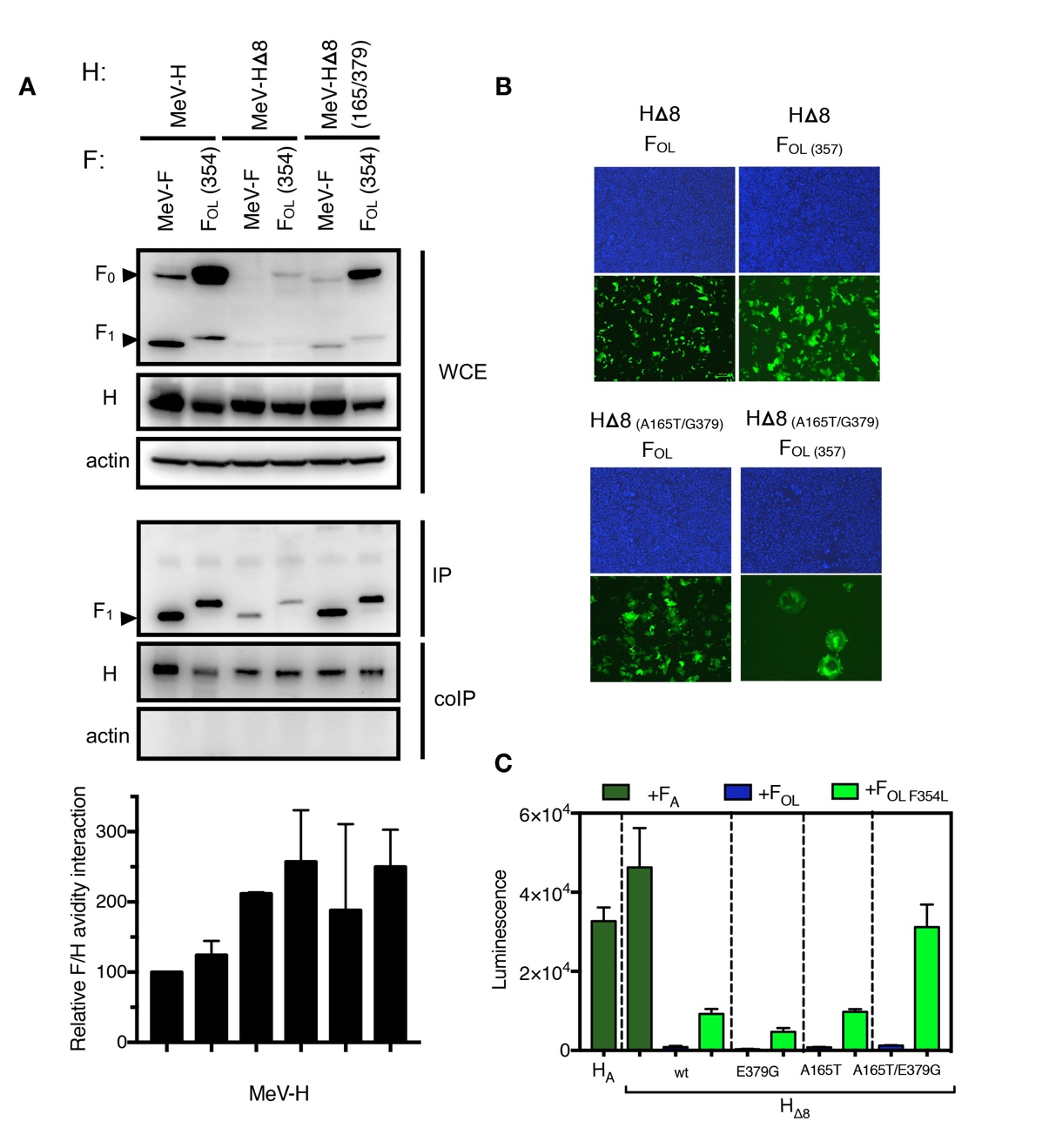

**Fig. S8.** Compensatory mutation for efficient rescue of the MR virus. (A) H/F coimmunoprecipitation. Transfected HEK293 cells were treated with the permeable cross-linker disuccinimidyl suberate (DSS) and subsequently lysed with radioimmunoprecipitation assay (RIPA) buffer. Complexes were then immunoprecipitated (IP) with anti-FLAG and protein G-Sepharose bead treatment. Proteins were boiled and subjected to immunoblotting using polyclonal anti-MeVH cyt to detect H antigenic material (co-IP). CoIP H proteins were detected in comparison with H proteins present in cell lysate prior to IP by immunoblotting using the same anti-H cyt antibody (WCE, whole cell extract). (B) Syncytium formation after cotransfection of Vero cells with different combination of H/F expression plasmids. A plasmid expressing EGFP was additionally used for easy visualization. 24 hours after co-transfection, cells were stained with 4’,6-diamidino-2-phenylindole (DAPI) and fluorescence in the appropriate channels was recorded. (C) Quantification of fusion activity for H/F pairs. A quantitative fusion assay was performed with baby hamster kidney (BHK) cells bearing the indicated H and F protein. 24 hours after transfection, cells were overlay onto Vero cells carrying the other half of the dual-split reported plasmid and luminescence was recorded 8 hours later.

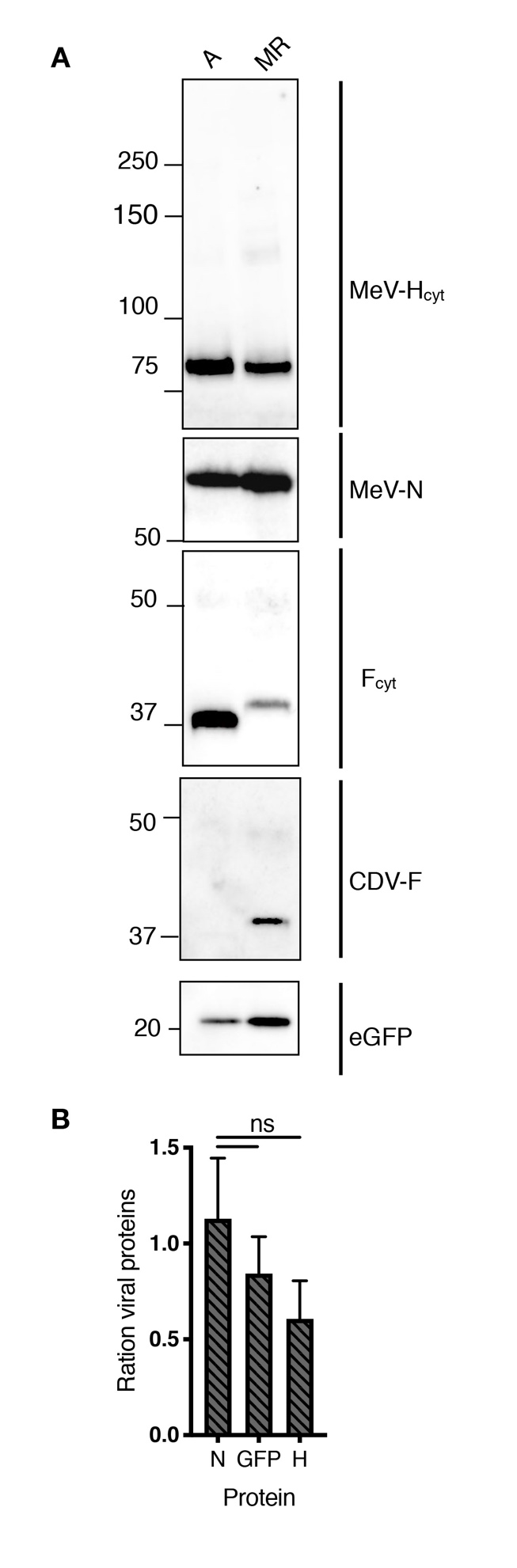

**Fig. S9**. Protein composition of the MR virus (A) Sucrose purified virus particles (1x10^5^ TCID_50_) were analyzed by Western Blotting using the indicated antibodies (B) Signal intensity from the bands were determined in 2 independent runs and plotted as the ratio of A vs MR virus. Ns, no significant (p=0.0.098), as determined by one-way ANOVA.

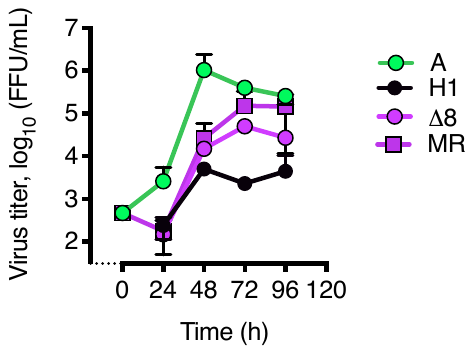

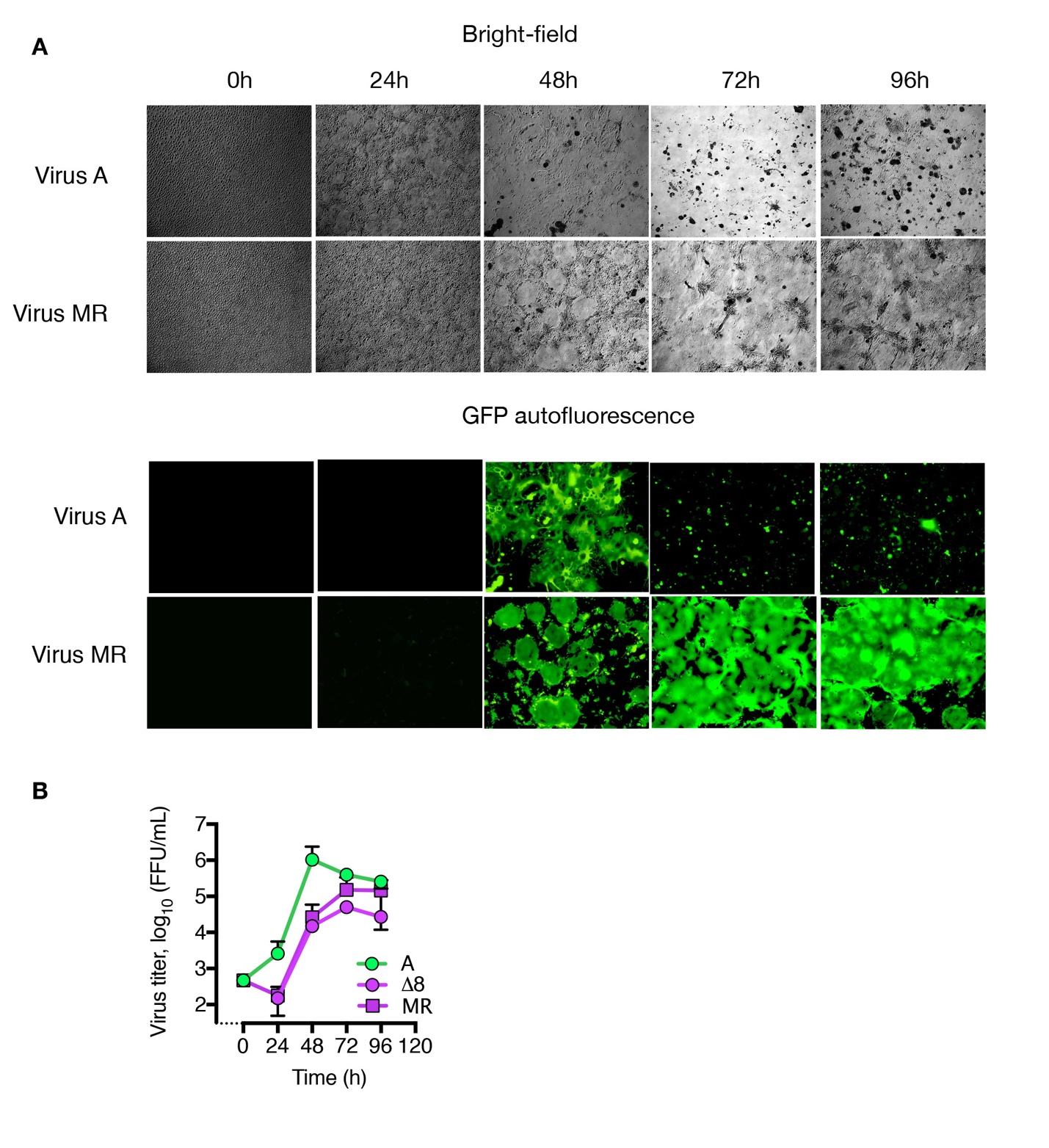

**Fig. S10.** Replication kinetics of the MR virus. (A) Replication of viruses on Vero/hSLAM infected at MOI of 0.03. At the indicated time, bright-field images and eGFP autofluorescence and were recorded. Magnification, ×40. (B) Infected cells were collected from the culture medium and virus titer was determined on Vero/hSLAM cells. MeV A, and H1, and Δ8 denote MeV expressing the corresponding MeV-H genes and MeV-F genotype A, whereas MR virus encodes MeV-H Δ8 plus CDV-F.

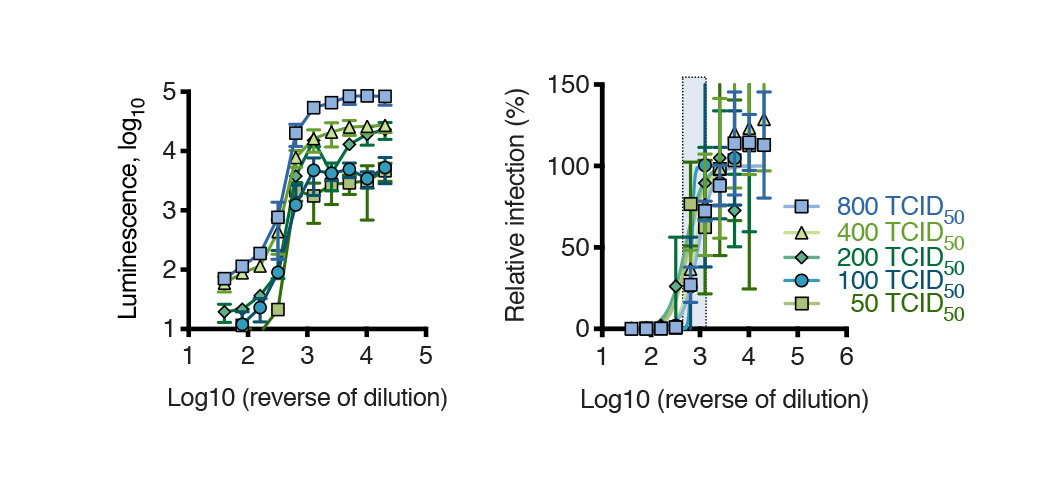

**Fig. S11**. Consistency of ND_50_ titers for human serum determined after infection of cells with MeV expressing firefly luciferase. A luciferase-based infection neutralization assay was performed using four different amounts of virus pre-incubated with serial dilutions of pooled human sera. Four days later, the luminesce (ie. infected-Vero cells) was determined quantitatively. Relative infection (percentage) was calculated by dividing measured luminescence in the presence of human serum by that the absence of human serum and converting to a percentage for each human serum dilution. Data points represent mean ± standard deviation of an experiment performed in quadruplicate. Data were fitted by non-linear regression analysis with Graph Pad Prism. The shaded area represents the dilution of human sera that inhibits virus infection of 50% (ND_50_). Regardless of the amount of virus input, ND_50_ remains constant.

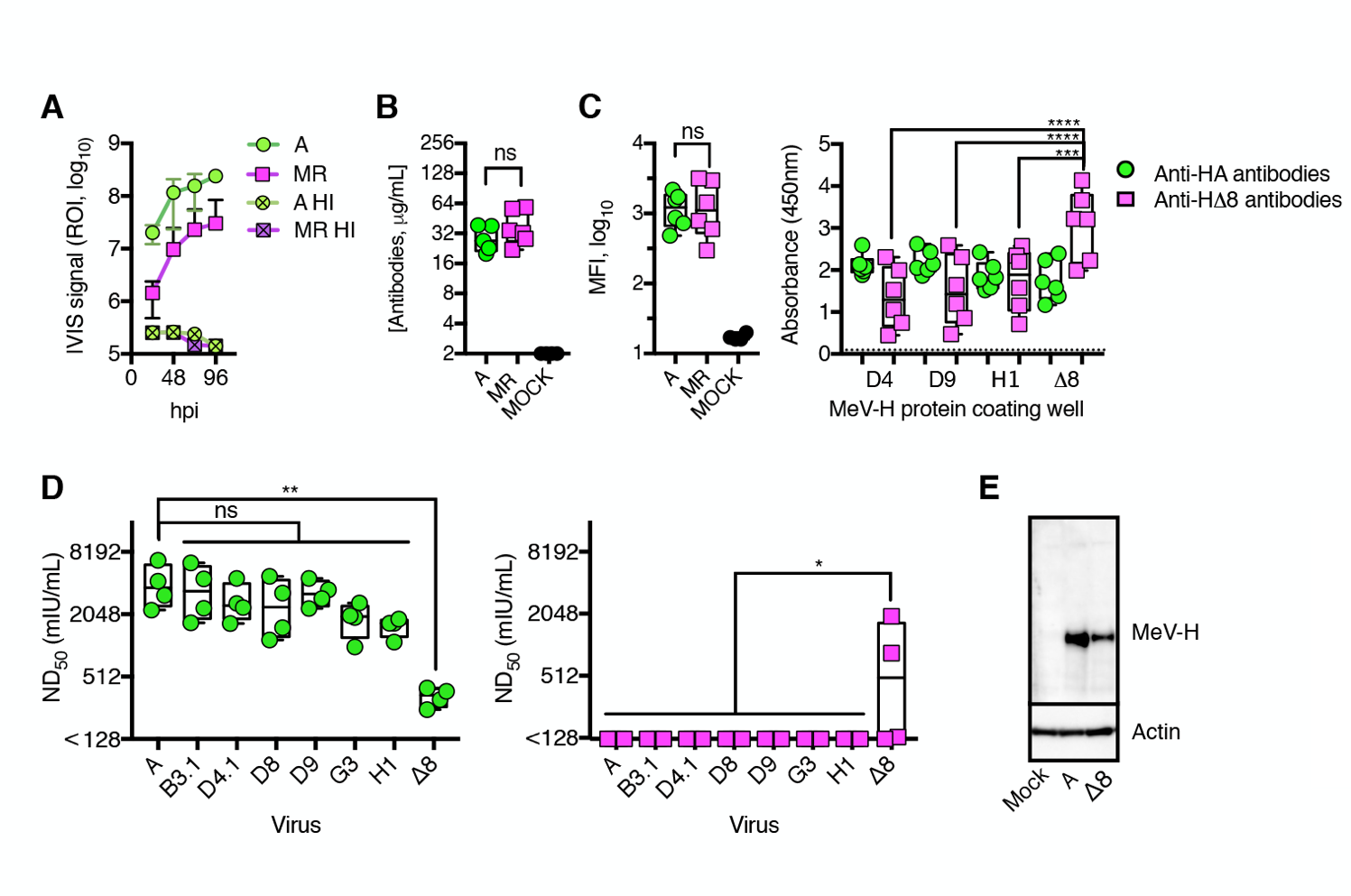

**Fig. S12.** Antibody Responses to Subdominant MeV-H Epitopes*.* MR virus infection in mice. (A) Replication of rMeV in HuCD46Ge-IFNar^KO^ mice. Groups of five mice were injected intraperitoneally with 2x10^5^ TCID_50_ particles that were either heat-inactivated (HI) or left untreated. Bioluminescence signal was measured at different time points using an *in vivo* imaging system (IVIS). (B) Antibody response towards the MeV-N protein of HuCD46Ge-IFNar^KO^ mice vaccinated with viruses. Groups of mice (n=6) were immunized intraperitoneally once with 2x10^5^ FFU particles. Antiserum samples were collected 4 weeks later, and anti-MeV-N antibodies were measured by enzyme-linked immunosorbent assay. Ns, no significant (p=0.2539), as determined by an unpaired two-tailed T test (t=1.219 with 9 degrees of freedom). (C), Antibody response to MeV-H from vaccinated mice. Left panel, MeV-H genotype A–specific IgG antibodies. Data were obtained by flow cytometry using transfected Mel-JuSo cells. Ns, no significant (p=0.6320) as per an unpaired two-tailed T test (t=0.194, with 10 degrees of freedom). Right panel, MeV-H genotype–specific response as determined by enzyme-linked immunosorbent assay using recombinant proteins. ****, p<.0001; ***, p=.0002, as determined by a two-way ANOVA with Turkey’s multiple comparison test. (D), DNA-immunization of mice with MeV-H A. Gene-based hydrodynamic injection of C57BL/6 mice was performed with MeV-H A (green) or MeV-H Δ8 (magenta). Neutralizing antibody response was next determined against currently circulating genotypes or mutant virus (Δ8) in serum samples of DNA-immunized mice. Significance was determined by Dunnett’s corrected 1-way ANOVA using the homotypic virus as reference. **, p=.0031; *, p<.02, with 24 degrees of freedom. (E) MeV-H expression from the plasmids used for DNA-immunization. Vero cells transfected with the indicated MeV-H were subjected to electrophoresis and immunoblotted with anti-MeV-Hcyt and β-actin (loading control).

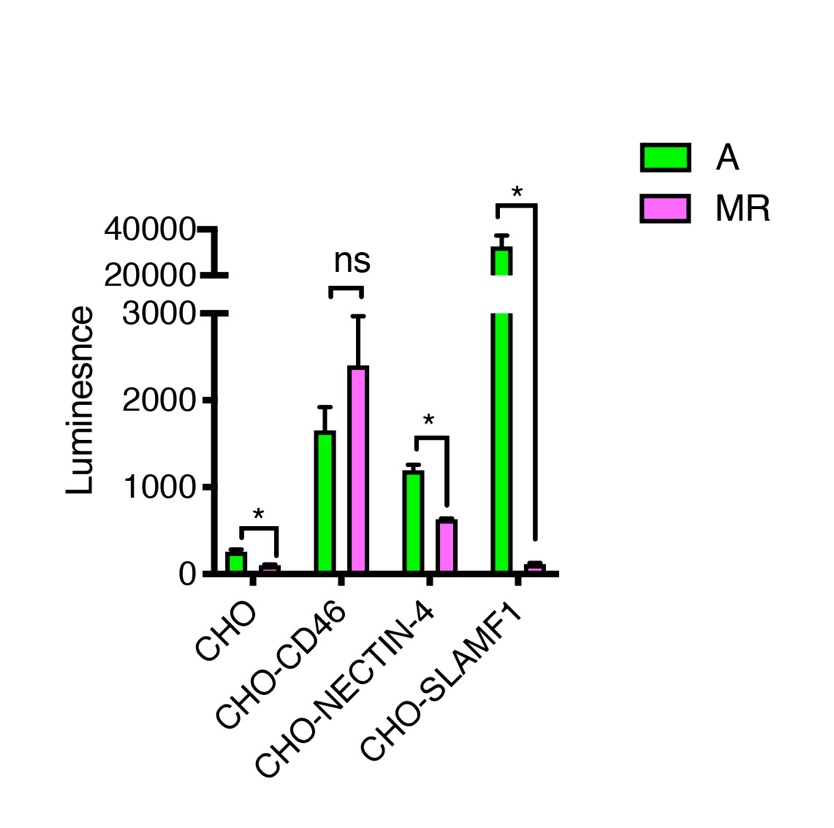

**Fig. S13**. Receptor-dependent infectivity of rMeV(Fluc)P. CHO cells expressing known MeV receptors were infected with the indicated virus at moi 0.1. Two-days after infection, luciferase signal was recorded. Statistical significance was determined using the T-test Holm-Sidak method with two degrees of freedom, with alpha=0.05.

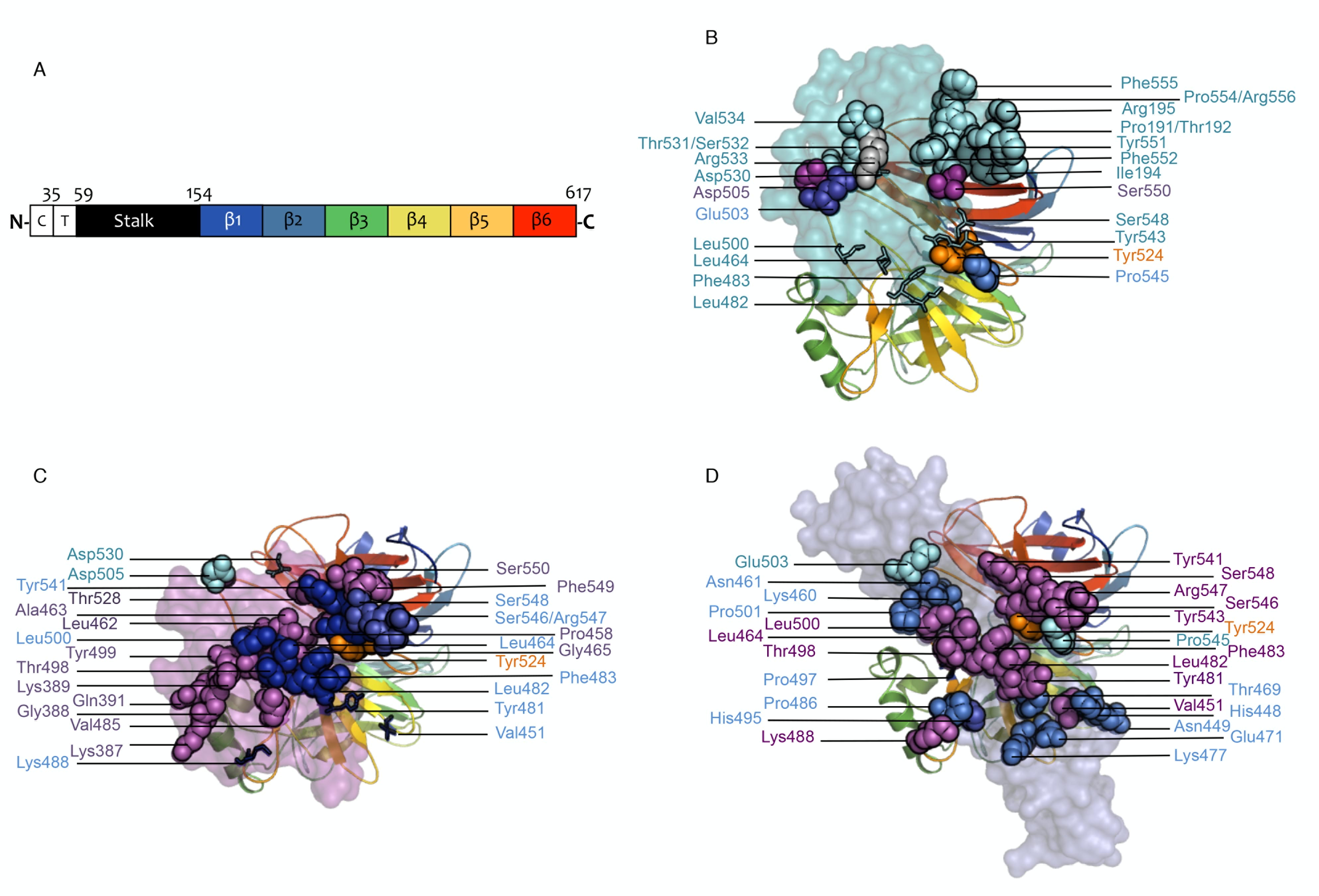

**Fig. S14.** Comparison of the receptors’ footprint in MeV-H. (A) Schematic representation of the MeV-H primary sequence. From left to right: C, cytoplasmic tail; T, transmembrane domain; Stalk domain; β1-6, beta-propeller blades 1-6. Amino acid position delineating the different domains are indicated. (B to D) MeV-H is shown as a ribbon with a rainbow coloring consistent with panel A, whereas the receptors are shown as translucent surface colored in cyan (SLAMF1, panel A), magenta (NECTIN4, panel B) and blue (CD46, panel C). Spheres denote residues ≤4.5Å from receptor entity SLAMF1 (PDB: 3ALZ), NECTIN4 (PDB: 4GJT) and CD46 (PDB: 3INB), colored in cyan, magenta and blue, respectively. In each MeV-H-receptor complex structure, spheres are colored differently to the receptor coloring if the residue is also relevant for the interaction with in any of the others receptors, and colored accordingly. Residue Y524 relevant for the interaction with all three receptors is colored in orange. For comparison, the MeV-H orientation is kept consistent across panels. Stick representation depict residues structurally involved as defined by ≤4.5Å from the receptor entity, but not functionally (L464, L482, F483, L500, D530, Y543, S548) and viceversa (V451, N-Y481, K488, P497) (*38*).

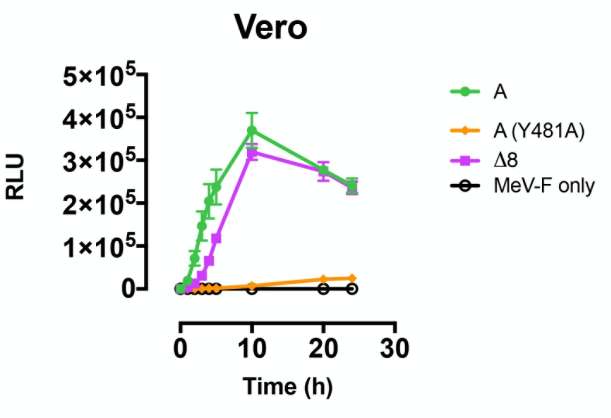

**Fig. S15.** Kinetic fusion assay on Vero cells. MeV-H vaccine strain or Δ8 were co-transfected together with MeV-F vaccine strain. Fusion was quantified by split-luciferase assay. Transfection of MeV-F alone served as negative control. Mean ± standard deviation of a representative experiment performed in triplicate.

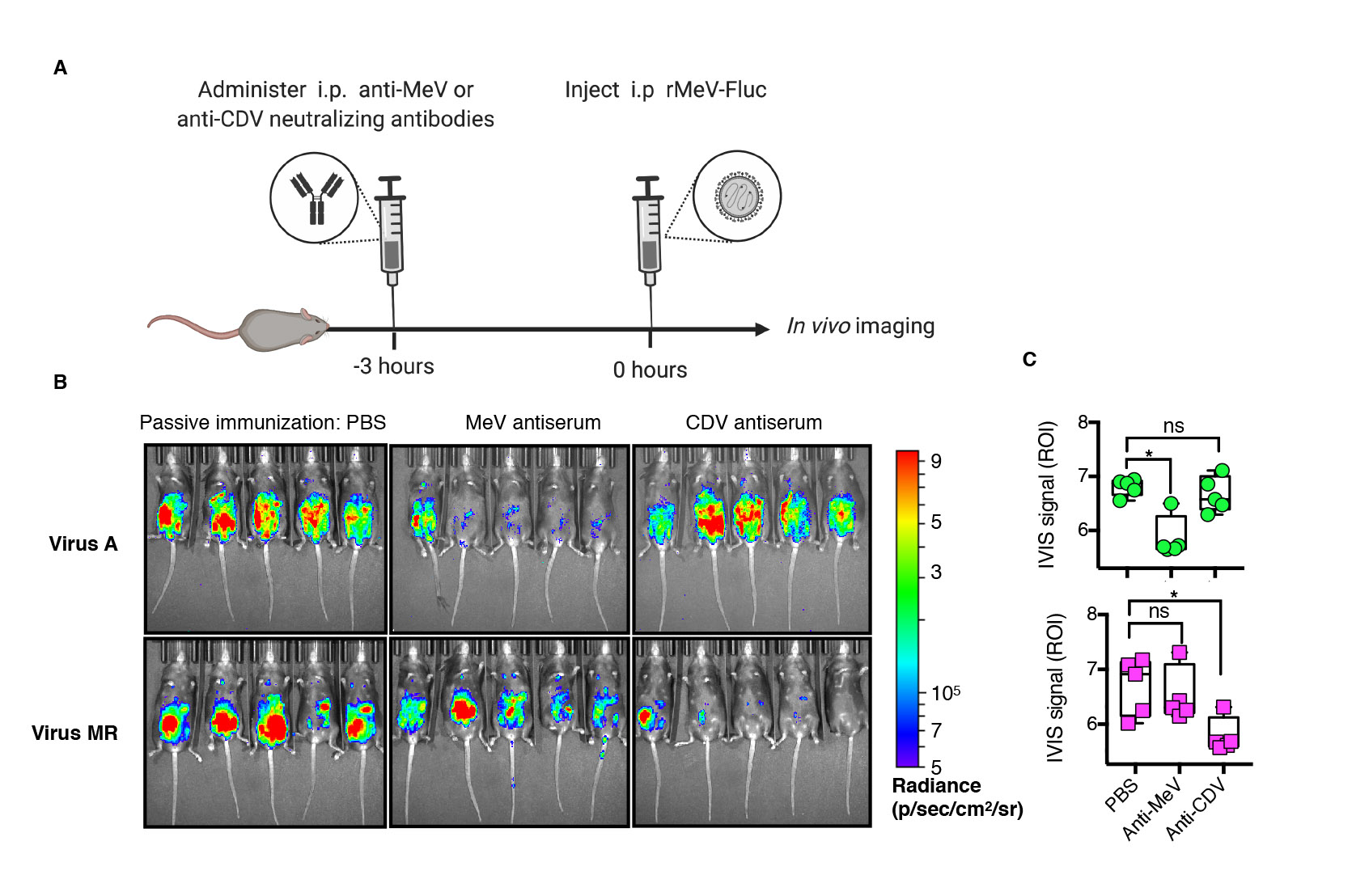

**Fig. S16.** The MR virus escapes *in vivo* neutralization by MeV-induced antibodies. (A) Passive immunization scheme. HuCD46Ge-IFNar^KO^ mice received 600 mIU of MeV or CDV polyclonal antiserum via the intraperitoneal route 3 hours before the injection of rMeV-expressing firefly luciferase (Fluc). In vivo imaging system (IVIS) was next used to detect bioluminescence signal, indicative of virus replication. B, Bioluminescence signal of virally encoded Fluc signal in mice pre-treated or not (PBS group) with anti-MeV or -CDV neutralizing antibodies-containing serum. The images were recorded 3 days after infection and they show absence of bioluminescence signal for MeV A in the presence of MeV antibodies whereas the presence of CDV antibodies, and not Me antibodies, inhibited the virus MR. (C) Quantification of the bioluminescence signal emitted from the abdominal and ventral area or region of interest (ROI), from each mouse infected with rMeV in the presence of passive immunity. PBS represents mock immunized mice. *, p<.005, as calculated with Dunnett’s corrected 1-way ANOVA with 12 degrees of freedom.

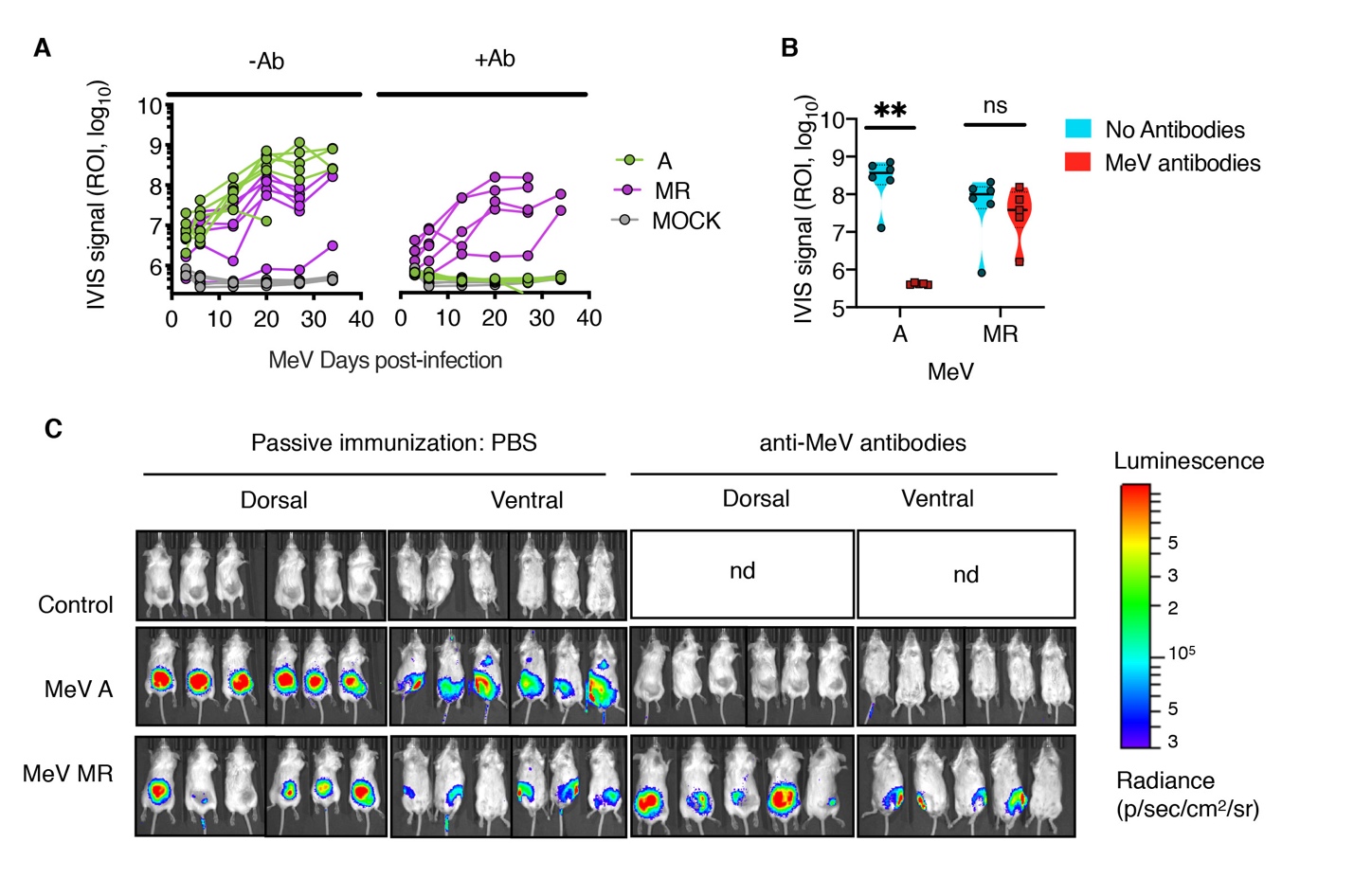

**Fig. S17.** MR trafficking to multiple myeloma tumors in the presence of anti-MeV antibodies. Mice bearing subcutaneous KAS 6/1 cells were treated intravenously with one dose of the indicated rMeV(Fluc) or PBS. Mice in the relevant group also received intraperitoneally 1000 mIU of anti-MeV antibodies three hours before MeV injecton. rin the presence or not and MeV-antibodies. (A) Bioluminescence quantitative signal (photons) was detected by *in vivo* imaging system (IVS) after intraperitoneal administration of 150mg/Kg of luciferin. (B) Comparative analysis of bioluminescence signal was determined at day 20 for MeV and MeV MR in the absecen or presence of anti-MeV antibodies. Nd, not done. **, p=0.0064, as determined by two-way ANOVA with Sidak’s multiple comparisons test. (C) Representative bioluminescence images of tumor-bearing SCID mice treated with MeV.

20.	O. T. Ertl, Tübingen, University, Diss., 2003., Erscheinungsort nicht ermittelbar (2003).
