## Supplementary material for "Pathogenic measles viruses cannot evolve to bypass vaccine-induced neutralizing antibodies": Peer Review File

#### Reviewer: 1

The authors generated a number of measles virus recombinants in which the known antigenic sites on the H protein have been modified so that the they become resistant to a panel of mouse monoclonal antibodies used in the past to define the 7 known epitope sites. They further identify a new 8th site and also that the F protein of measles virus has epitopes for neutralising antibodies. In order to generate a virus that would allow them to study the effect of sera on measles H protein they therefore had to make a further recombinant virus in which the F gene of measles virus is replaced with that of CDV. With this virus they carry out some mouse experiment showing that this virus (MR) can replicate in a mouse treated passively treated with a measles serum. In general, I have only minor comments and suggestions on small inaccuracies on the vast number of experiments which are detailed below.

The main questions that the paper raised are:

1 The authors refer to the intramural discussions on the DURC aspects of the study. Figure 3 gives a reassurance that the MR cannot infect the various CHO recombinant cells expressing the three known measles receptors. An extra experiment involving the use of the MeV #8 virus would have been helpful to ascertain that the possibly suboptimal interaction between the F protein of CDV and the H protein of MeV with modifications in the globular head region but not in the stalk part of the H protein, was also not able to grow in the various CHO cells.

2 The study is of course limited to epitopes recognized by the mouse. The question as to what extent mouse data may pertain to human immunity is one that should be discussed. Several supplementaries deal with this question but it may be worthwhile to discuss this question.

3 Similarly, the question of applicability to human immunity, the authors use in the IVIS experiment in supplementary S13 a guinea pig serum rather than a human serum. Might it be worthwhile to include such an experiment?

4 A further question that is potentially of relevance to providing reassurance on DURC aspects is the potential that there may be neutralising epitopeson non-envelope proteins of the virus. Nobody has understood the remarkable and early response to the N protein in human sera.

Minor comments and suggestions:

1 Line 56 “vaccines” should be “vaccinees”.

2 In line 135 why do the authors refer to 5 sites when their data purport to indicate that even one of the 7 sites is sufficient to retain the monotypic nature of the virus though assessed against a rabbit serum in S4.

3 Line 232 I think the authors should justify the use of the EB V antibodies to standardize the dilutions of sera

4 Line 494 “syncytium” rather than “syncytia”.

5 Line 469 should MeV-F (condition 3 be condition 2)?

6 Line 452 E471E should that be E471K?

7 In Figure S 14 there is reference to the various #7 mutants with one of the site intact. Although digging deep in the past literature published by this group, one may be able to find the correct sequences of these I would suggest a Table of the mutations that were present in each of the H proteins se virus would be helpful as a supplementary.

### Reviewer: 2

##### (Cross-Review)

#### Reviewer: 2

The authors have generated recombinant MeVs in a background of an attenuated strain, introduced H and F glycoproteins of wild-type MeV and/or CDV and systematically modified the glycoproteins to modulate VN epitopes. By screening for serum neutralization, they identified the existence of several antigenic sites in H, of which some were shown to be critical for binding to CD150 or nectin-4. On basis of these data they conclude that evolution of pathogenic MeVs capable of escaping serum neutralization in vaccinated individuals is nearly impossible.

The approach used in this study seems counter-intuitive: the authors use a vaccine strain as background, introduce the H of a wild-type strain but modify this to bind CD46. The rationale for this (instead of using the native H of vaccine strain Moraten) is not well-explained. The introduction of the three mutations required to generate tropism for CD46 may also have an influence on specific VN epitopes?

Although the manuscript contains interesting data, the inability of MeV to escape VN antibodies has already been well-explained by the group of Makoto Takeda by demonstrating that the receptor-binding region of H represents an important VN epitope (Tahara et al., JVI 2013, not cited in the manuscript). Subsequently the group of Benhur Lee has used insertional mutagenesis to demonstrate biological constraints on the genetic and antigenic variability of MeV, especially in the transmembrane glycoproteins (Fulton et al 2015, Beaty et al 2016). Therefore, I do not believe that this manuscript meets the requirements for publication in Science.

Specific comments

1. Line 12: the cited reference does not fit with the statement in this sentence. The current reference 5 is more suitable.

2. Line 13: eradication is not (yet) a formal goal of the WHO, at this moment the organization aims for regional elimination and mortality reduction.

3. Line 14/15: formal receptor names are CD150 and nectin-4. The authors are encouraged to use CD150 instead of SLAM in figures (eg Fig 3, S1, S6) and text (eg Vero-hCD150 instead of Vero-hSLAM) and write nectin-4 without capitals.

4. Line 17-19: the MeV polymerase has a high mutation rate, but MeV does not. Measles genome sequences are remarkably stable, and show virtually no drift during outbreaks. This sentence could easily be misinterpreted by readers with limited background knowledge of MeV.

5. Lines 21-24: Tahara et al were the first to demonstrate that the receptor-binding site of the MeV H constitutes a conserved neutralizing epitope (JVI 2013). This observation should be acknowledged, and the original publication should be cited.

6. Line 24: consider adding Beaty et al (Viruses 2016)

7. Line 29, reference 10: consider using Greenwood et al, Vaccine 2016

8. Line 38: H1 is a genotype, not a strain.

9. Line 40, Fig S1: DNA transfection panels have limited added value over infection panels.

10. Line 42. Fig S2: the statement “Of note, C2 viruses selectively escape neutralization” should be modified to include “…by this mAb” to avoid misinterpretation.

11. Line 43, Fig S3: I do not agree with the statement “Neutralization assays confirmed that viruses encoding the MeV-H Δ8 escaped all 30 mAbs”. Several mAbs (eg BH068, BH021, BH101) show substantial inhibition, although not complete neutralization at the dose range used.

12. Line 48: Fig 1A does not show VN titer of mouse anti-H antiserum

13. Line 50: change “all other” to “the other four”

14. Lines 52-54: it is not clear to me how the authors came to this conclusion, I do not see it supported by the data.

15. Line 56: I assume that by ‘serum from measles vaccines’ the authors mean polyclonal sera from vaccinated humans?

16. Line 60: a seven-fold decrease in VN titer against the delta-8 virus seems modest as compared to the results obtained with mAbs. Apparently polyclonal VN Abs show more plasticity? It is unclear to me why the VN data are plotted against Epstein-Barr VCA IgG and not serum dilution. This is also not explained in the materials and methods section in the supplemental data.

17. Line 67-72: MeV-d7-III seems to interact less efficient with particularly nectin4 (but also CD46), is this important? Are other antigenic sites (besides III) also involved in receptor binding?

18. Line 78-79: did the authors check the effect of introduction of these compensatory mutations in H on immunogenicity?

19. Line 79-81: There is no need for Fig 2 to be a main figure, this can easily be supplemental, as it does not contribute to the main message of the manuscript but is purely method description. Panel B should include all three viruses. In panel C, it is unclear why growth curves were determined in Vero-hCD150 cells (viruses do not bind CD150?). In panel D, it should explicitly be stated if the ‘human samples’ are from vaccinate or naturally infected subjects. The authors describe a left panel, but there is no right panel.

20. Line 94-95: did the authors test these sera for cross-reactivity with CDV? Is it possible that some human sera contain antibodies that bind to CDV F?

21. Line 99 + 101: it is not clear why the authors use the term ‘weakly’. No significant differences between A and MR mice in Fig S10.

22. Lines 104-122: it is not clear to me why exclusive CD46 receptor usage is so important? It does not support the main message of the manuscript, and therefore it is questionable whether this should be in the main figures.

23. Conclusion 4 was published in 2013 (Tahara et al JVI, see above).

#### Reviewer: 3

**(Cross-Review)**

Overall, I confirm my initial recommendation. This is a good work, but not good enough for Science. I share the views of Reviewer 2. Perhaps it could be a good paper for Science Translational Medicine.

#### Reviewer: 3

This paper reports a heroic effort to finally prove the well-known and obvious observation that the measles virus is not able to escape vaccine immunity. The authors show that vaccination raises neutralizing antibodies against five independent epitopes. Since mutating less than five provides no selective advantage for the virus, it is impossible for the virus to escape all of them at the same time.

Briefly, the authors developed a panel of measles viruses (MeV) mutants with selective B cell neutralizing epitopes knock-outs in the viral hemagglutinin protein. At least five epitopes have to be knocked out to abrogate polyclonal neutralization. Viruses that accumulate mutations and escape neutralization become inefficient in binding human receptors, thus concluding that the probability of emergence of a neutralization-resistant phenotype is close to zero.

The present work dives into the pathophysiology of measles virus elucidating mechanisms of infectivity and immune evasion with elegant in vitro studies. However, despite the vaccine strain remains serologically monotypic, short duration of protection, decreased vaccine coverage and appearance of other MeV genotypes could pose an increasing threat. Currently, there is no evidence about whether MeV is escaping vaccine-induce immunity. The findings of the present work could partially answer to these questions, but they are narrow and do not really advance the research of better solutions for measles.

Overall, I am not sure this work is a manuscript for Science.

### Reviewer: 4

##### (Cross-Review)

The comments of the other reviewers are helpful. They reinforce my general impression that the main interest to the general reader in the manuscript lies in the provocative – but not fully supported – claim contained within the title. That said there are clearly new and important data in the present study, and I note that none of the reviewers have serious reservations about the experiments. A more judiciously worded manuscript could still be of interest.

#### Reviewer: 4

Measles has a high mutation rate but a vaccine isolated in 1954 still confers full protection. The authors offer an explanation for this phenomenon, that antibodies to each of eight co-immunodominant sites on MeV haemagglutinin are individually capable of neutralisation, and that these include the receptor binding surface for SLAMF1 and NECTIN4. This is inferred from introduction of mutated haemagglutinins to the Moraten vaccine strain. They argue that mutations would have to arise in multiple sites simultaneously for measles to evade vaccine-induced immunity, and that this is mathematically implausible given the 1/10^5 mutation rate of the RdRP. They further contend that even if such an unlikely event were to occur, this could only be accomplished by losing affinity for the pathogenicity determining receptors.

There is a weaknesses in this argument: it cannot be assumed that the average mutation rate of 1/10^5/site/round of replication is uniform. What is the mathematical distribution of mutation rates for individual polymerases? I'm not sure we know this, I'd be surprised if it were a normal distribution with a low standard deviation. Rare events might introduce multiple mutations simultaneously. For example, there could be 'mutator' polymerases arising themselves by mutation that – although not generally favourable for the virus – might nevertheless produce progeny that transiently accumulated a high number of mutations. Furthermore the virus might acquire a high number of focussed mutations from encounter with a host mutator, for example adenosine deaminase enzymes. It is also possible that measles could acquire new properties by recombination. Similarly, although the authors demonstrate that the receptor binding site for SLAMF1 and NECTIN4 is a target for neutralisation (antigenic site III, figure S6), it's not impossible that measles could acquire a new tropism by binding to a completely different receptor that conferred pathogenicity and transmissibility.

In general the experiments are conducted well and the data are convincing. There's not any obvious way I can think of that the authors could address these points experimentally. Whereas determining the statistical distribution of error rates for the measles polymerase would be interesting it's clearly outside the scope of this manuscript. I would prefer that the authors address these points in discussion as caveats to their conclusion.

Minor comments:

Figure 1C is a bit small, if the data points could be spaced out further it would make it easier to appreciate the point.

Figure 2A – are these sucrose gradient purified virions?

Figure S1 – it would be good to see expression levels of the receptors (as per figure 3B) and the transfected mutant MeV-Hs (as per figure S14?). It's quite a busy figure, could the authors summarise the findings in an accompanying table?

Figure S3 – could the authors indicate why the X axes are variable between experiments?

Figure S6 – again, could the expression levels be indicated as per figure 3B?

#### Reviewer: 1

The authors generated a number of measles virus recombinants in which the known antigenic sites on the H protein have been modified so that the they become resistant to a panel of mouse monoclonal antibodies used in the past to define the 7 known epitope sites. They further identify a new 8th site and also that the F protein of measles virus has epitopes for neutralising antibodies. In order to generate a virus that would allow them to study the effect of sera on measles H protein they therefore had to make a further recombinant virus in which the F gene of measles virus is replaced with that of CDV. With this virus they carry out some mouse experiment showing that this virus (MR) can replicate in a mouse treated passively treated with a measles serum. In general, I have only minor comments and suggestions on small inaccuracies on the vast number of experiments which are detailed below.

We would like to thank this reviewer for taking the time to carefully review our manuscript and for the positive input.

The main questions that the paper raised are:

1 The authors refer to the intramural discussions on the DURC aspects of the study. Figure 3 gives a reassurance that the MR cannot infect the various CHO recombinant cells expressing the three known measles receptors. An extra experiment involving the use of the MeV #8 virus would have been helpful to ascertain that the possibly suboptimal interaction between the F protein of CDV and the H protein of MeV with modifications in the globular head region but not in the stalk part of the H protein, was also not able to grow in the various CHO cells.

The experiment proposed by this reviewer was indeed performed and is shown in Figure 3A. The MeV MR is the one which includes the multiply mutated measles virus hemagglutinin (MeV-H) plus the canine distemper virus fusion protein (CDV-F). Given the importance of this result we have now included additional quantitative data for the infectivity of this virus on the various

CHO cells via luciferase measurement using versions of the MeV A and MeV MR viruses into which we engineered a luciferase transgene (Fig. S13).

2 The study is of course limited to epitopes recognized by the mouse. The question as to what extent mouse data may pertain to human immunity is one that should be discussed. Several supplementaries deal with this question but it may be worthwhile to discuss this question.

Neutralization by human sera is the major focus of our studies. See for example Figure 1C in which human sera were depleted of F-specific antibodies and used to neutralize viruses encoding the unmodified (A) or multiply mutated (Δ8) MeV H protein. However, in light of the reviewer suggestion we have now dedicated Figures 2 and 3 exclusively to the quantitative analysis of MeV A and MeV MR neutralization activity of human serum samples. Data using measles specific mouse (Fig. 1B), rabbit (new Fig. S5) and guinea Pig (new fig. S16 and S17) antisera are still included in the manuscript and support previous conclusion by Yewdell's lab studying influenza virus, where antibodies from mice, guinea pigs, chickens and lamprey were found to target similar antigenic sites (Cell Host Microbe 2013; 13:314-323; Nature 2014; 430: 174-180; Nature Immunology 2017; 18;456-463). We previously showed that the antigenic site III (which overlaps the receptor-binding site) is targeted both by convalescent sera and by serum from vaccine recipients (Virus Res 2017; 236: 30-43).

Similarly, the question of applicability to human immunity, the authors use in the IVIS experiment in supplementary S13 a guinea pig serum rather than a human serum. Might it be worthwhile to include such an experiment?

We agree with the reviewer. However, the reason we used guinea pig serum in this study was that, based on pilot studies, we could only observe statistical reduction in luminescence signal when mice were passively immunized with ≥ 600 mIU. In light of the IACUC-imposed limit to the allowable volume of serum that could be transferred for passive immunization, and the antibody titers of the human sera that were available to us, the experiment was not practicable using human serum. Hence we used a guinea pig serum known to contain a very high titer of MeV neutralizing antibodies.

To further convince this reviewer of the antibody evasion by the MR virus, we have included as a new Fig. S16, a comparative study of virus trafficking to multiple myeloma tumor xenografts in the presence or not of anti-MeV-antibodies. Since the virus was administered intravenously and the tumor was engrafted subcutaneously, we obtained a more convincing demonstration of the effect of passive immunization. In this human xenograft model, the virus MR trafficked to the tumor site despite the presence and neutralizing antibodies, whereas the Moraten vaccine could not.

4 A further question that is potentially of relevance to providing reassurance on DURC aspects is the potential that there may be neutralising epitopes on non-envelope proteins of the virus. Nobody has understood the remarkable and early response to the N protein in human sera.

We appreciate the interesting suggestion. We concur that anti-N antibodies may impede measles pathogenesis even though they lack virus neutralizing activity. For example, it has been reported that the N protein has immunosuppressive properties via inhibition of B cell antibody production (Ravanel et al., 1997. Measles virus nucleocapsid protein binds to FcgRII and inhibits human B cell antibody production. J Exp Med 186: 269-278). However, we think that this speculation would be beyond the scope of the current manuscript.

Minor comments and suggestions:

1 Line 56 “vaccines” should be “vaccinees”.

Corrected

2 In line 135 why do the authors refer to 5 sites when their data purport to indicate that even one of the 7 sites is sufficient to retain the monotypic nature of the virus though assessed against a rabbit serum in S4.

The data in Figure S4 (now S5) do indeed show that only two of the eight antigenic sites in Δ8 could be mutated without impacting neutralization resistance to polyclonal rabbit antiserum. However, the data in Figure 1 B show that this applies to three of the eight sites when using polyclonal mouse antiserum. We therefore refer to the more conservative estimate of 5 codominant antigenic sites.

3 Line 232 I think the authors should justify the use of the EB V antibodies to standardize the dilutions of sera

Thanks for pointing this out. We have modified the sentence to read as: "Epstein-Barr virus viral capsid antigen (VCA) IgG levels were used to accounting for dilution factors since depletion conditions should not affect their levels"

4 Line 494 "syncytium" rather than "syncytia".
Corrected

5 Line 469 should MeV-F (condition 3 be condition 2)?
Thanks for pointing out an error with the conditions. Indeed, the mistake laid on condition 1. The second time it is mentioned should have said condition 2 (MeV-H depletion) since condition 1 represents incubation with untransfected cells.

6 Line 452 E471E should that be E471K?
Corrected

7 In Figure S 14 there is reference to the various #7 mutants with one of the site intact. Although digging deep in the past literature published by this group, one may be able to find the correct sequences of these I would suggest a Table of the mutations that were present in each of the H proteins se virus would be helpful as a supplementary.

Thanks for the suggestion. We have added a Table of mutations as a supplementary.

Reviewer: 2

(Cross-Review)

#### Reviewer: 2

The authors have generated recombinant MeVs in a background of an attenuated strain, introduced H and F glycoproteins of wild-type MeV and/or CDV and systematically modified the glycoproteins to modulate VN epitopes. By screening for serum neutralization, they identified the existence of several antigenic sites in H, of which some were shown to be critical for binding to CD150 or nectin-4. On basis of these data they conclude that evolution of pathogenic MeVs capable of escaping serum neutralization in vaccinated individuals is nearly impossible.

We would like to emphasize that we have shown that the multiple antigenic sites are CODOMINANT.  Hence, stepwise mutation of those sites does not provide the virus with an evolutionary pathway to generate a new serotype. Instead, all codominant sites must be simultaneously lost to impact the ability of the virus to escape antibody neutralization, which is the main determinant of resistance to measles infection.

The approach used in this study seems counter-intuitive: the authors use a vaccine strain as background, introduce the H of a wild-type strain but modify this to bind CD46. The rationale for this (instead of using the native H of vaccine strain Moraten) is not well explained.

We thank the reviewer for the opportunity to clarify this point. Our experimental design was to incorporate a series of selected spontaneous monoclonal nAb escape mutations into the MeV H glycoprotein. We previously noted that MeV genotype H1 carries one of the most antigenically advanced of naturally occurring MeV-H proteins, having already become resistant to several of the nAb known to neutralize the genotype A (Edmonston) [Muñoz-Alía et al., 2013. PLos One 13(2): e52306]. Choosing the H protein from MeV genotype H1 as a starting point enabled us to minimize alterations of the MeV-H structure which had previously been reported to be rigid and intolerant of mutations [Fulton BO et al., 2015. Cell Rep 11(9); 1331-8]. The text referenced this rationale as “we introduced increasing numbers and varying combinations of epitope escape mutations into an antigenically advanced and relatively malleable MeV-H protein obtained from the H1 genotype of MeV, which to facilitate the evaluation of mutants with disrupted SLAMF1 and NECTIN-4 receptor binding sites…"

The introduction of the three mutations required to generate tropism for CD46 may also have an influence on specific VN epitopes?

We did consider this question but were unable to demonstrate an effect of these mutations on the dominant VN epitopes. Figure 1B shows efficient serum neutralization of multiple MeVs displaying H proteins from various non-CD46 tropic MeV genotypes, but greatly reduced neutralization of the Δ8 virus which remains CD46 tropic. These data do not provide evidence to support the idea that new epitopes are created by mutations conferring CD46 tropism. Moreover, the three mutations that we introduced to generate CD46 tropism did not impact susceptibility to the individual monoclonal antibodies in our panel and the neutralization sensitivity of the original H1-derived CD46-tropic virus was indistinguishable from the parental H1 virus. CD46 tropism was therefore retained in all of the mutant viruses that we subsequently evaluated and our finding confirm that CD46 tropism can be disregarded as a significant factor in antibody-neutralization escape.

Although the manuscript contains interesting data, the inability of MeV to escape VN antibodies has already been well-explained by the group of Makoto Takeda by demonstrating that the receptor-binding region of H represents an important VN epitope (Tahara et al., JVI 2013, not cited in the manuscript). Subsequently the group of Benhur Lee has used insertional mutagenesis to demonstrate biological constraints on the genetic and antigenic variability of MeV, especially in the transmembrane glycoproteins (Fulton et al 2015, Beaty et al 2016). Therefore, I do not believe that this manuscript meets the requirements for publication in Science.

These authoritative comments argue that our work lacks novelty in light of two published manuscripts.

On the first point, that the inability of measles to escape antibody neutralization has been already well-explained, the Tahara et al., 2013 paper cited by this reviewer reported a monoclonal antibody (2F4) that could block receptor attachment sites of the measles H glycoprotein. The authors showed that this antibody epitope mapped into the receptor binding domain (RBD) and hyphothesized that since the virus would need to mutate its RBD to escape neutralization, this might represent significant evolutionary constraint. However, in the same manuscript the authors described single point mutations in the H glycoprotein that circumvented the neutralizing activity of the 2F4 antibody and restored the interactions with SLAMF1 or NECTIN4.

Indeed, there is an interesting history around the 2F4 epitope described by Tahara *et al*. The authors acknowledge in the paper that 2F4 recognizes an epitope previously described to be associated with other already reported mAbs. In particular, mAb cl55 (available at the European Cell Collection Bank, Cat# 95040311) was shown to bind R533 and impact receptor binding back in 2004 (Massé et al., JVI 2004). When the structure of the MeV-H in complex with the receptors become available, one of the authors of the current manuscript studied potential overlap in footprints between the SLAMF1 and CD46 receptors (Please see Fig. 3 at Santiago et al., 2010. Nat Struc Mol Bio 17(1); 124-9) leading to the suggestion that CD46 usage could have arisen as a strategy to circunvent neutralization. This led to the mapping of certain neutralizing antibody epitopes to the receptor-binding footprint and provided evidence that certain clinical isolates present mutations in the RBS that impact monoclonal antibody binding (Muñoz-Alía et al., 2017. Virus Research 236: 30-43). It was therefore suggested that immune escape might have to associate with a lack of SLAMF1 binding which would make it theoretically impossible to generate escape mutants.

In the current manuscript, we have shown that the virus can indeed mutate the RBS and escape neutralization by antibodies targeting this antigenic site, but that this is accomplished via the use of alternative receptos. However, our current work shows that a far more important constraint to the antigenic evolution of the virus is the existence of multiple nonoverlapping codominant epitopes in the envelope. We further proved the contribution of the F protein to the monotypic nature of measles virus (Figures 1C and new Figure 2). In summary, our findings provide a new model for the monotypic nature of the virus.

Since the Tahara study did not evaluate or demonstrate the dominance or codominance of multiple antigenic sites on the H protein, it did not elucidate (as the current study does) why measles cannot escape from a polyclonal antibody response. Regarding the suggestion to include this Tahara reference in the manuscript, we already included a reference to M. Tahara *et al.*, *Measles Virus Hemagglutinin Protein Epitopes: The Basis of Antigenic Stability*. *Viruses* **8**, (2016)), a review article which has the same first and last authors and which includes additional data over Tahara et al, JVI, 2013.

As to the second point of this reviewer, that our manuscript lacks novelty or significance in light of Fulton et al 2015 and Beaty et al 2016, we believe that this reviewer comment further strengthens the argument that our manuscript sheds important new light on the monotypic nature of measles and should therefore be published in Science to help address a widely held misconception in the measles field, namely that measles virus is monotypic because its H and F proteins are "structurally inflexible".

The group of Benhur Lee in their original article (Fulton BO et al., 2015; Cell Rep 11: 13331-1138) and subsequent review article (Beaty et al 2016) claimed to have discovered another “factor constraining antigenic variability of measles virus”. Fulton et al., 2015 describes a study using shotgun transposon mutagenesis to demonstrate that the measles F and H proteins are less tolerant of peptide insertions compared to other proteins encoded in the measles genome. However, the group of Roberto Cattaneo (Navaratnarajah CK et al., 2011. Nat Struct Mol Biol. 18: 128-134) was able to successful perform insertional mutagenesis in at least 6 strategic locations, which retargeted the virus to use a pseudoreceptor. None of these locations were identified in the transposon mutagenesis studies of Fulton et al., 2015. Further, in the current manuscript we found that it was not difficult to construct and rescue numerous recombinant measles viruses with multiple mutations in their H proteins. Indeed, one of our reported constructs contains as many as 25 amino acid changes scattered throughout the H protein when compared with the closest known MeV-H (genotype H1). Further, we were able to generate replication competent viruses in which we completely replaced the measles F protein with the CDV F protein without destroying virus infectivity. Thus, although not stated in the manuscript, our findings and the findings of others do not support the conclusion of the Fulton paper, that measles evolution is constrained by the inelasticity of its surface glycoproteins, a conclusion that has become generally enrooted in the field, as the review's comments indicate.

In view of the current COVID19 pandemic and the role of individual amino acids in the SARS-CoV-2 spike on antigenicity (Li etl., 2020; Cell, doi: 10.1016/j.cell.2020.07.012) together with recent reports of antigenic variation in mumps virus (Rasheed et al., 2019; PNAS116(38); 19071-76) we believe our work sheds important insights into the mechanism that truly constrains the serotypic evolution of measles virus.

Specific comments

1. Line 12: the cited reference does not fit with the statement in this sentence. The current reference 5 is more suitable.

We have included both references in line 12 now since we believe that Mina et al. (ref.1) elegantly showed the extent of immune suppression induced by measles virus infection.

2. Line 13: eradication is not (yet) a formal goal of the WHO, at this moment the organization aims for regional elimination and mortality reduction.

Thanks for pointing this out. "Eradication" has been substituted by "elimination"

3. Line 14/15: formal receptor names are CD150 and nectin-4. The authors are encouraged to use CD150 instead of SLAM in figures (eg Fig 3, S1, S6) and text (eg VerohCD150 instead of Vero-hSLAM) and write nectin-4 without capitals.

It is not clear to us how this reviewer draws such a conclusion since searches for CD150 are automatically -redirected to the pertinent sites (https://www.ncbi.nlm.nih.gov/gene/?term=CD150). Since we are referring to human genes/

proteins, we capitalized the names accordingly, following the guidelines by the HUGO Gene Nomenclature Committee, then avoiding confusion with the homologous receptors from other paramyxoviruses.

4. Line 17-19: the MeV polymerase has a high mutation rate, but MeV does not. Measles genome sequences are remarkably stable, and show virtually no drift during outbreaks. This sentence could easily be misinterpreted by readers with limited background knowledge of MeV.

The text has been updated

5. Lines 21-24: Tahara et al were the first to demonstrate that the receptor-binding site of the MeV H constitutes a conserved neutralizing epitope (JVI 2013). This observation should be acknowledged, and the original publication should be cited.

Please see our extensive discussion of the Tahara et al. reference above. We included a review article by the same authors instead of referring to this paper, since they expanded their previous observations in the follow up review article. As to who was the first to demonstrate that the receptor-binding site of the MeV-H constitutes a conserved neutralizing epitope, this epitope was first reported as III (see new table1) by Sheshberadaran and Norbby, 1986; Virology 152; 58-54), IIIA/IIIB by Hummel and Bellini, 1995; JVI 69: 1913-1916) and E2 by our group (Lech et al., 2013. PLOS One 8:e52306). We refer this reviewer to Santiago et al., 2010. (Fig.3; Nat Struc Mol Bio 17(1); 124-9) for further clarification on the overlap between receptor footprint and neutralizing antibody epitopes.

6. Line 24: consider adding Beaty et al (Viruses 2016)

This is a review article about "factors that constraint the monotypic nature of the virus" and as suggested above by the reviewer, we included the original publication instead (Fulton et al, 2015). Please be aware of space constraints with the Science Report format.

7. Line 29, reference 10: consider using Greenwood et al, Vaccine 2016

Thanks for the suggestion, the reference has been updated.

8. Line 38: H1 is a genotype, not a strain.

We have updated the text.

9. Line 40, Fig S1: DNA transfection panels have limited added value over infection panels.

We have eliminated DNA transfection from Fig. S1.

10. Line 42. Fig S2: the statement "Of note, C2 viruses selectively escape neutralization" should be modified to include "…by this mAb" to avoid misinterpretation.

We apologize for the inaccuracy; the text has been modified.

11. Line 43, Fig S3: I do not agree with the statement "Neutralization assays confirmed that viruses encoding the MeV-H Δ8 escaped all 30 mAbs". Several mAbs (eg BH068, BH021, BH101) show substantial inhibition, although not complete neutralization at the dose range used.

Our statement is based on the fact that 4-fold or greater reduction in serum neutralization is used to define antigenic variation in Influenza Virus (Smith et al., 2004; Science 305(5682): 371-376) or serotype distinction in dengue virus (Katzelnick et al., 2015; Science 349(6254): 1338-1343). However, to satisfy this reviewer concern, we have revised the text to read "Neutralization assays confirmed that viruses encoding the MeV-H Δ8 showed a decrease in neutralization susceptibility to all 30 mAbs".

12. Line 48: Fig 1A does not show VN titer of mouse anti-H antiserum

We apologize for the typo, we now refer to Fig. 1B instead.

13. Line 50: change "all other" to "the other four"

Corrected

14. Lines 52-54: it is not clear to me how the authors came to this conclusion, I do not see it supported by the data.

With the exception of antigenic site IIB, elimination of antigenic sites gave identical results, in terms of a significant antigenic different with both mouse (Figure 1B) and rabbit (Figure S5) measles virus-antiserum.

15. Line 56: I assume that by 'serum from measles vaccines' the authors mean polyclonal sera from vaccinated humans?

Corrected

16. Line 60: a seven-fold decrease in VN titer against the delta-8 virus seems modest as compared to the results obtained with mAbs. Apparently polyclonal VN Abs show more plasticity? It is unclear to me why the VN data are plotted against Epstein-Barr VCA IgG and not serum dilution. This is also not explained in the materials and methods section in the supplemental data.

We agree with the reviewer. We believe that there is an additive effect of antibodies involved in neutralization in polyclonal preparations. Also, in addition to the codominant epitopes we have identified, our data point to the existence of multiple subdominant B cell epitopes in the H glycoprotein to explain the residual susceptibility to neutralization of the MR virus, thus providing an additional constraint to virus evolution

On the reviewers second point, we have updated the text to clarify the use of Epstein-Barr VCA IgG: "Epstein-Barr virus viral capsid antigen (VCA) IgG levels were used to accounting for dilution factors since depletion conditions should not affect their levels". We believe this method accounts more realistically for different dilutions introduced into the human serum due to the use of different conditions.

17. Line 67-72: MeV-d7-III seems to interact less efficient with particularly nectin4 (but also CD46), is this important? Are other antigenic sites (besides III) also involved in receptor

binding?

We thank the reviewer for pointing out this interesting finding. Since none of the reported co-crystal structures could satisfactorily explain our results, we believe different mutations in the antigenic site might have allosteric interactions and this a matter of investigation. Indeed, different MeV-H from different genotypes display noticeable differences in binding to the receptors (https://repositorio.uam.es/handle/10486/662949). We alluded to this fact in the text as: "We identified no second-site mutations in known contact residues to explain this unexpected segregation of CD46 and NECTIN4 tropisms and therefore postulate that the phenotype may be partially attributable to specific noncontact residues in the MeV-H protein of genotype H1".

18. Line 78-79: did the authors check the effect of introduction of these compensatory mutations in H on immunogenicity?

We did not observe any effect on antigenicity for the compensatory mutations.

19. Line 79-81: There is no need for Fig 2 to be a main figure, this can easily be supplemental, as it does not contribute to the main message of the manuscript but is purely method description. Panel B should include all three viruses. In panel C, it is unclear why growth curves were determined in Vero-hCD150 cells (viruses do not bind CD150?). In panel D, it should explicitly be stated if the 'human samples' are from vaccinate or naturally infected subjects. The authors describe a left panel, but there is no right panel.

Following the reviewer suggestion, we have divided Figure 2 between Figure 3 and supplementary figures (new Fig. S9 and S10). Vero-hCD150 cells were used because we were comparing the growth kinetics of MeV A, MeV Δ8, MeV MR and MeV H1. MeV H1 uses CD150 as a receptor and cannot use CD46. Hence Vero-hCD150 were used to allow the amplification of all four viruses. Data from the group of Dr. Yanagi (PMID:12050387) showed that the titer of a recombinant virus expressing MeV-H A was the same in Vero and Vero-CD150. However, we observed a much lower titer for MeV H1, which we attributed to the extensive cytopathic effect via the use of CD150 in this virus and we considered that its replication capacity could not be assessed properly. Given this reviewer demand, this growth curve is now shown.

As to the origin of the serum human samples, this is detailed as follows in Materials and Methods: "Human serum samples were from 3 cohorts. Cohort 1 samples were obtained from the Erasmus Medical Center serum bank and have been previously described [9]. Members of cohort 1 likely were never exposed to wild-type MeV and likely received a monovalent measles vaccination at age 14 months and a measles-mumps-rubella (MMR) vaccination at age 9 years. Cohort 2 samples were obtained from the Mayo Clinic Biobank in Olmsted Country (Rochester, MN); this region has no documented evidence of circulating measles or rubella virus in the past and has high vaccine coverage. However, MMR vaccination status was not available for most samples. The subjects' age for cohort 2 ranged from 30 to 44 years, and 80% were female. Cohort 3 samples were custom purchased from Innovative Research Inc, based on age and race. Hispanic donors between 20 and 29 years old were chosen to increase the likehood of obtaining serum samples from vaccinated people."

20. Line 94-95: did the authors test these sera for cross-reactivity with CDV? Is it possible that some human sera contain antibodies that bind to CDV F?
Indeed, we tended to observed a more significant antibody evasion when we used the MeV Δ 8 and human sera depleted from MeV-F antibodies. Given that there is a 67% amino acid homology between CDV and MeV F proteins, there might exist cross-neutralizing antibodies that decrease our significance with MeV MR.

21. Line 99 + 101: it is not clear why the authors use the term 'weakly'. No significant differences between A and MR mice in Fig S10.
The ELISA results did not have sufficient power to detect statistically significant differences due to the high dispersion in the values. However, it can be observed that some samples did not react at all with the different hemagglutinins and we believe this is an important finding that needed to be included. We have added two panels to the new Fig. S12 where mice immunized with the

MeV-H Δ 8 protein can only neutralize the homotypic virus, and not other measles virus genotypes.

22. Lines 104-122: it is not clear to me why exclusive CD46 receptor usage is so important? It does not support the main message of the manuscript, and therefore it is questionable whether this should be in the main figures.

CD46 receptor usage was critical to the completion of the study since it allowed us to evaluate the impact of epitope ablating mutations that destroy the binding of the virus to SLAMF1 and to NECTIN-4. Since our findings contradict previous science that showed that CD46 and NECTIN-4 tropism are indivisible (Mateo M. et al., 2013; JVI 87(16):9208-16 and Liu Y-P. et al., 2014; JVI 88(4):2195-204), we believe that our data supporting only one tropism should be highlighted.

23. Conclusion 4 was published in 2013 (Tahara et al JVI, see above).

Again, we refer to our earlier discussion of this point. Sheshberadaran and Norbby, 1986 (Virology 152; 58-54), Hummel and Bellini, 1995 (JVI 69: 1913-1916), Massé et al., 2004 (JVI 78(17): 9051-9063), Santiago et al., 2010. (Nat Struc Mol Bio 17(1); 124-9) and Muñoz-Alía et al., 2017. (Virus Research 236: 30-43) have presented data showing that the RBD is a neutralizing epitope. Please see also Table 1. Here, we reconfirm this result and further demonstrate that, when the virus is adapted to use alternative receptor (CD46), the true extent of its inability to circumvent a polyclonal antibody response becomes fully apparent.

#### Reviewer: 3

(Cross-Review)

Overall, I confirm my initial recommendation. This is a good work, but not good enough for Science. I share the views of Reviewer 2. Perhaps it could be a good paper for Science Translational Medicine.

We thank the reviewer for this recommendation. We would like to refer this reviewer to the responses we have provided above to fully address the concerns expressed by reviewer 2, particularly those regarding the novelty and significance of the work.

#### Reviewer: 3

This paper reports a heroic effort to finally prove the well-known and obvious observation that the measles virus is not able to escape vaccine immunity. The authors show that vaccination raises neutralizing antibodies against five independent epitopes. Since mutating less than five provides no selective advantage for the virus, it is impossible for the virus to escape all of them at the same time.

Briefly, the authors developed a panel of measles viruses (MeV) mutants with selective B cell neutralizing epitopes knock-outs in the viral hemagglutinin protein. At least five epitopes have to be knocked out to abrogate polyclonal neutralization. Viruses that accumulate mutations and escape neutralization become inefficient in binding human receptors, thus concluding that the probability of emergence of a neutralization-resistant phenotype is close to zero.

The present work dives into the pathophysiology of measles virus elucidating mechanisms of infectivity and immune evasion with elegant in vitro studies. However, despite the vaccine strain remains serologically monotypic, short duration of protection, decreased vaccine coverage and appearance of other MeV genotypes could pose an increasing threat. Currently, there is no evidence about whether MeV is escaping vaccine-induce immunity. The findings of the present work could partially answer to these questions, but they are narrow and do not really advance the research of better solutions for measles. Overall, I am not sure this work is a manuscript for Science.

We thank this reviewer for taking his time to review our work. The author states that "short duration of protection, decreased vaccine coverage and appearance of other MeV genotypes" could pose an increasing threat and that our work could partially answer those questions. While we agree that decreased vaccine coverage is a problem, that matter is beyond the scope of our manuscript.  We do not agree that short duration of protection or the appearance of other MeV genotypes is an issue for measles since childhood vaccination generally leads to lifelong seropositivity and protection from the disease. The question our paper addresses is to explain why measles virus remains monotypic and cannot escape the protective effect of vaccination, even though the vaccine was developed 70 years ago. A literature search indicates that the monotypic nature of the measles virus is still under debate: Fatemi Nasab et al., 2016. Pathog Dis 74(8); 089; Kneissl et., 2012. PLos One 7(10);e46667; Lévy etl., 2012. Mol Ther 20(9): 1699-712; Kweder et., 2014. Adv Virol. 205617. For a recent article on this topic we refer this reviewer to an article recently published by Michael Mina's group "Measles vaccine immune escape; should we be concerned" Eur. J Epidemiol 34(10): 893-896.

The reviewer is also concerned that our findings are "narrow and do not really advance the research of better solutions for measles" In contrast, we believe our findings do have far reaching implications for measles control:

First, by demonstrating that the virus cannot evolve to escape the polyclonal antibody response that follows measles vaccination, our work gives strong support to the idea that it may be possible to eradicate measles. Only by elucidating the basis of its monotypic nature, as we have done in this manuscript, can it be appreciated that the probability of an evolutionary pathway to generate an escape mutant is vanishingly small.

Birrer et al. first published in *Nature* the existence of measles antigenic variants by using monoclonal antibodies against the hemagglutinin (Birrer et al., 1981. Nature 293;67-69). But despite the high mutation rate of its polymerase, the virus remains monotypic and there have been several attempts to explain this, for example on the basis of the proximity of H protein receptor-binding and neutralizing antibody-targeted motifs (Hashiguchi et al., 2007. PNAS 104; 19535-19540) or the lack of plasticity of the viral coat proteins (Fulton et al., 2015. Cell Rep. 11: 13331-1338). However, we here show that all 8 antigenic sites on the measles virus hemagglutinin must be ablated to allow the virus to escape just the H-glycoprotein-directed polyclonal response. We further show that both measles virus H and F antibodies are responsible for protection. Measles virus Hemagglutinin protein has long been considered the main mediator of protection in vaccinated or pre-exposed individuals (de Swart et al., 2009. JGV 90:2982-2989; de Swart et al., 2005. JGV 79:11547-11551). As a result, measles serologic status are based only on antibody responses to the H protein and essentially ignore the antibody response to the F protein which we now show is equally effective and therefore highly significant as an additional measure of virus protection.

Although not discussed in the manuscript, a second area where our work may help to advance the search for better measles solutions is that the antibody resistant MR virus we describe in the paper could provide the basis for a vaccine effective in the presence of maternal antibodies. A major barrier to measles eradication has to do with the timing of measles vaccination. Newborn children are generally protected from measles by antibodies that they acquire transplacentally from their mothers in utero (Niewiesk, 2014. Front Immunol 5:446). Unfortunately, in addition to being resistant to measles infection, the babies of measles-immune mothers are also resistant to measles vaccination because their transplacentally acquired antibodies neutralize the vaccine. For this reason, vaccination is not initiated until 12 to 15 months of age in countries with low measles incidence versus 9 to 12 months of age in countries with higher incidence (Orenstein et al., 1986. Dev Biol Stand 65:13-21). In this regard, our finding that the measles virus vaccine can be engineered so as to create a derivative virus that is not only effective in the presence of antibodies from vaccinated mothers, but which also retains the ability to stimulate the production of antibodies is of evident interest at a moment. This possibility is being pursued but is beyond the scope of the current manuscript.

#### Reviewer: 4

(Cross-Review)
The comments of the other reviewers are helpful. They reinforce my general impression that the main interest to the general reader in the manuscript lies in the provocative – but not fully supported – claim contained within the title. That said there are clearly new and important data in the present study, and I note that none of the reviewers have serious reservations about the experiments. A more judiciously worded manuscript could still be of interest.

We thank the reviewer for this insight and agree that none of the reviewers have serious reservations about the experiments we conducted.

#### Reviewer: 4

Measles has a high mutation rate but a vaccine isolated in 1954 still confers full protection. The authors offer an explanation for this phenomenon, that antibodies to each of eight coimmunodominant sites on MeV haemagglutinin are individually capable of neutralisation, and that these include the receptor binding surface for SLAMF1 and NECTIN4. This is inferred from introduction of mutated haemagglutinins to the Moraten vaccine strain. They argue that mutations would have to arise in multiple sites simultaneously for measles to evade vaccine-induced immunity, and that this is mathematically implausible given the 1/10^5 mutation rate of the RdRP. They further contend that even if such an unlikely event were to occur, this could only be accomplished by losing affinity for the pathogenicity determining receptors.

Thanks to the reviewer for this cogent summary.

There is a weaknesses in this argument: it cannot be assumed that the average mutation rate of 1/10^5/site/round of replication is uniform. What is the mathematical distribution of mutation rates for individual polymerases? I'm not sure we know this, I'd be surprised if it were a normal distribution with a low standard deviation. Rare events might introduce multiple mutations simultaneously. For example, there could be 'mutator' polymerases arising themselves by mutation that – although not generally favourable for the virus – might nevertheless produce progeny that transiently accumulated a high number of mutations. Furthermore the virus might acquire a high number of focussed mutations from encounter with a host mutator, for example adenosine deaminase enzymes. It is also possible that measles could acquire new properties by recombination. Similarly, although the authors demonstrate that the receptor binding site for SLAMF1 and NECTIN4 is a target for neutralisation (antigenic site III, figure S6), it's not impossible that measles could acquire a new tropism by binding to a completely different receptor that conferred pathogenicity and transmissibility.

In general the experiments are conducted well and the data are convincing. There's not any obvious way I can think of that the authors could address these points experimentally. Whereas determining the statistical distribution of error rates for the measles polymerase would be interesting it's clearly outside the scope of this manuscript. I would prefer that the authors address these points in discussion as caveats to their conclusion.

We would like to thank the reviewer for his/her valuable comments and appreciate this input.

The suggestion that mutations may be specifically targeted to the regions of the H protein that need to be mutated in order to escape antibody neutralization is interesting, but lacks supporting data. Indeed, the fact that antibody escape mutants of MeV have not arising in 70 years of measles vaccination suggests that there may be no such targeting. Whereas mutators may initially benefit from low fidelity polymerases, they lose fitness against antimutators and are eventually extinguished. This selection against mutation load should result in mutation rates being pushed even lower. Contrary to the reviewer suggestion ADAR is anticipated to have little to no effect on mutation rate on the envelope because it preferentially edits defective measles virus genomes. Regular genomes are encapsidated and cannot therefore be efficiently edited (Pfaller CK et al., 2018; PLos Biology 16(11); e2006577). Another important factor relates to the different cell lines and whether their mutation rates would be different, Interestingly, the VSV mutation rate in different cell lines was determined by Combe and Sanjuan 2014, (PLos Pathogens10(1):e1003855), and shown to be invariable despite the wide variety of different cell lines, including those with p53 inactivated. We therefore believe our calculations provide a good estimate.

Minor comments:

Figure 1C is a bit small, if the data points could be spaced out further it would make it easier to appreciate the point.

We have made Fig1C a slightly wider. We hope this helps.

Figure 2A – are these sucrose gradient purified virions?

Yes, they are. We apologize for this inadvertent omission; gradient purified virion will not show F0 but the cleaved product. We have updated the text to indicate this.

Figure S1 – it would be good to see expression levels of the receptors (as per figure 3B) and the transfected mutant MeV-Hs (as per figure S14?). It's quite a busy figure, could the authors summarise the findings in an accompanying table?

We appreciate this reviewer's suggestion. However, based on the strong recommendation of reviewer#2, we have eliminated DNA-transfection from the figure since we agree that cell-to-cell fusion might not be the same as virus envelope fusion. We have now included expression levels of the receptors in Fig. S1B.

Figure S3 – could the authors indicate why the X axes are variable between experiments?
Please notice that some graphs show concentration in ng/mL whereas other panels show dilution instead.

Figure S6 – again, could the expression levels be indicated as per figure 3B?
Does this reviewer suggest to copy the data included now in Fig. S1B (the expression levels would be the same)?
